# Specialized Tfh cell subsets driving type-1 and type-2 humoral responses in lymphoid tissue

**DOI:** 10.1101/2022.07.28.501817

**Authors:** Saumya Kumar, Afonso P. Basto, Filipa Ribeiro, Silvia C. P. Almeida, Patricia Campos, Carina Peres, Sarwah Al-Khalidi, Anna Kilbey, Jimena Tosello, Eliane Piaggio, Momtchilo Russo, Margarida Gama-Carvalho, Seth B. Coffelt, Ed W. Roberts, Helena Florindo, Luis Graca

**Affiliations:** Instituto de Medicina Molecular, Faculdade de Medicina, Universidade de Lisboa, 1649-028 Lisboa, Portugal; Instituto Gulbenkian de Ciência, Oeiras, Portugal; CIISA - Centro de Investigação Interdisciplinar em Sanidade Animal, Faculdade de Medicina Veterinária, Universidade de Lisboa, 1300-477 Lisboa, Portugal; Laboratório Associado para Ciência Animal e Veterinária (AL4AnimalS), Lisbon, Portugal; Research Institute for Medicines (iMed.ULisboa), Faculty of Pharmacy, Universidade de Lisboa, Av. Prof. Gama Pinto, 1649-003 Lisboa, Portugal; Institute of Cancer Sciences, University of Glasgow, Glasgow, UK; Cancer Research UK Beatson Institute, Glasgow, UK; Institut Curie, PSL Research University, INSERM U932, F-75005 Paris, France; Institute of Biomedical Sciences, Department of Immunology, University of Sao Paulo, Sao Paulo 05508-000, Brazil; BioISI – Biosystems & Integrative Sciences Institute, Faculty of Sciences, University of Lisboa, 1749-016 Lisboa, Portugal

**Keywords:** Tfh cell, germinal center, isotype switching, single-cell transcriptomics, bioinformatics

## Abstract

Effective antibody responses are essential to generate protective humoral immunity. Different inflammatory signals polarize T cells towards an appropriate effector phenotype during an infection or immunization. Th1 and Th2 cells have been associated with the polarization of humoral responses for several decades. However, it is now established that T follicular helper cells (Tfh) have a unique ability to access the B cell follicle and support the Germinal Centre (GCs) responses by providing help to B cells. We investigated the specialization of Tfh cells induced under type-1 and type-2 conditions. We first studied homogenous Tfh cell populations generated by adoptively transferred TCR-transgenic T cells in mice immunized with type-1 and type-2 adjuvants. Using a machine learning approach, we established a gene expression signature that discriminates Tfh cells polarized towards type-1 and type-2 response, defined as Tfh1 and Tfh2 cells. The Tfh1 and Tfh2 distinct signature was validated against datasets of Tfh cells induced following LCMV or helminth infection. Using single-cell transcriptomics, we also dissected the heterogeneity of Tfh cells from the two immunizing conditions. Our results show that Tfh cells acquire a specialized function under distinct types of immune responses, but with the coexistence of a small population of Tfh cells of the alternative type. Furthermore, the specific molecular hallmarks of Tfh1 and Tfh2 cells identified herein offer putative new targets for tuning humoral responses.

## Introduction

Humoral immunity plays a central role in protective responses against infection as well as in pathological responses in allergy and autoimmunity. The formation of Germinal Centres (GCs) is the main event underlying the production of high-affinity antibodies essential for protective (or pathogenic) immunity (*1*). A landmark finding in the history of immunology was the notion that B cells require help from CD4 T cells for class switching and affinity maturation (*2, 3*). This finding led to the designation of CD4 T cells as helper T cells. However, the generalization of helper function to all CD4 T cells was challenged with the identification of a subset of CD4 T cells, the T follicular helper (Tfh) cells, with the unique ability to access B cell follicles and provide help to B cells (*4–8*). The differentiation of Tfh cells was found to be dependent on the transcription factor Bcl6 (*9–11*). Although most studies on Tfh cells elucidated the overall Tfh function regarding their interactions with B cells within GCs, few studies have looked into the distinct functional subsets of Tfh cells required for appropriate antibody responses to different types of immunization or infection (*12–14*).

Another historical advance in immunology was the discovery of effector CD4 T cell specialization, based on their inflammatory milieu, towards a Th1 or Th2 phenotype (*15, 16*). This finding led to a better understanding of the characteristics of an immune challenge in the selection of adequate effector mechanisms against different pathogens. An established paradigm in this specialization is the selection of humoral responses leading to type-1 or type-2 antibody production (namely IgG2a versus IgG1/IgE isotypes in mice) following infection by viruses (such as LCMV) or parasites (such as helminths) (*17*). The polarized effector CD4 T cells involved in type-1 and −2 responses, Th1 and Th2 cells, have been well studied, having a well-defined cytokine profile and a characteristic transcriptional regulation. Among their most distinctive features, Th1 cells are characterized by *T-BET* expression and production of IFNγ, while Th2 cells express *GATA3* and produce IL-4, IL-5, and IL-13 (*18*). However, Th1 and Th2 cells do not directly promote GC responses or engage GC B cells. These characteristics are unique to Tfh cells (*3*). Thus, Tfh cells are likely driving affinity maturation and isotype switching under type-1 or type-2 responses. Recent findings showed the production of type-specific cytokines, IFN-γ and IL-4, by Tfh cells, suggesting their specialization (*12, 14, 19*). However, the biology of such specialized Tfh subsets remains poorly defined. Indeed, unlike Th1 and Th2 cell polarization, Tfh cell polarization is difficult to study *in vitro* due to the requirement of multiple cellular interactions with distinct cell types (*20*). Furthermore, the heterogeneous nature of *in vivo* immune responses induced with type-1 or −2 pathogens, as there may not be a “pure” type-1 or type-2 response, also creates difficulties (*21, 22*).

We circumvented the obstacles to studying Tfh polarization with the combination of two approaches. First, we designed near homogenous *in vivo* conditions to generate controlled humoral responses, allowing the comparison of the transcriptome of Tfh cells from polarized conditions. For that, we combined the transference of TCR-transgenic T cells, the use of adjuvants inducing clearly polarized type-1 or −2 humoral responses, and immunized with a defined target antigen (ovalbumin) without additional proteins. We used two different strains of mice (BALB/c and C57Bl/6), known to be more prone to type-2 and −1 polarization, respectively, to gain greater power to identify a transcriptional signature independent of strain bias. A machine learning approach allowed the deduction of a transcriptional signature for type-1 or type-2 Tfh cells that we validated with cells from public datasets of LCMV and helminth infections.

Second, to investigate the heterogeneity of type-1 and −2 polarized Tfh subsets, we generated single-cell transcriptomes from mice immunized with the polarizing adjuvants. The single-cell datasets allowed dissection of Tfh subset heterogeneity within both immunizations. We found a minor subpopulation of Tfh2 cells in mice subjected to type-1 immunization, and a minor subpopulation of Tfh1 cells following type-2 immunization. These minor populations of divergent Tfh cells are coherent with the detection of a small amount of immunoglobulins of divergent types following immunization with the two adjuvants.

Using these combined approaches, we were able to describe the polarized Tfh subsets generated under type-1 and −2 immune responses and dissect the heterogeneity within each Tfh subpopulation. These results elucidate the biology of Tfh cell subsets arising following type-1 and −2 polarization and shed light on the specialized Tfh-B cell help in GC responses.

## Results

### Generation of Tfh cells under type-1 and type-2 immune responses

We first defined the appropriate adjuvants to bias the immune response towards type-1 and type-2 conditions, using IgG1 and IgG2a as surrogate markers of type-2 or type-1 responses in mice (*23*). We immunized C57BL/6 mice with ovalbumin (OVA) using incomplete Freund’s adjuvant (IFA), CpG-oligodeoxynucleotides (CpG), or nanoparticles containing OVA and CpG (NP-CpG) as the adjuvant (**Fig. 1A**). We found that immunization with OVA-IFA could reliably lead to OVA-specific IgG1, while CpG or NP-CpG promoted the production of OVA-specific IgG2a antibodies (**Fig. 1B**).

**Figure 1.**
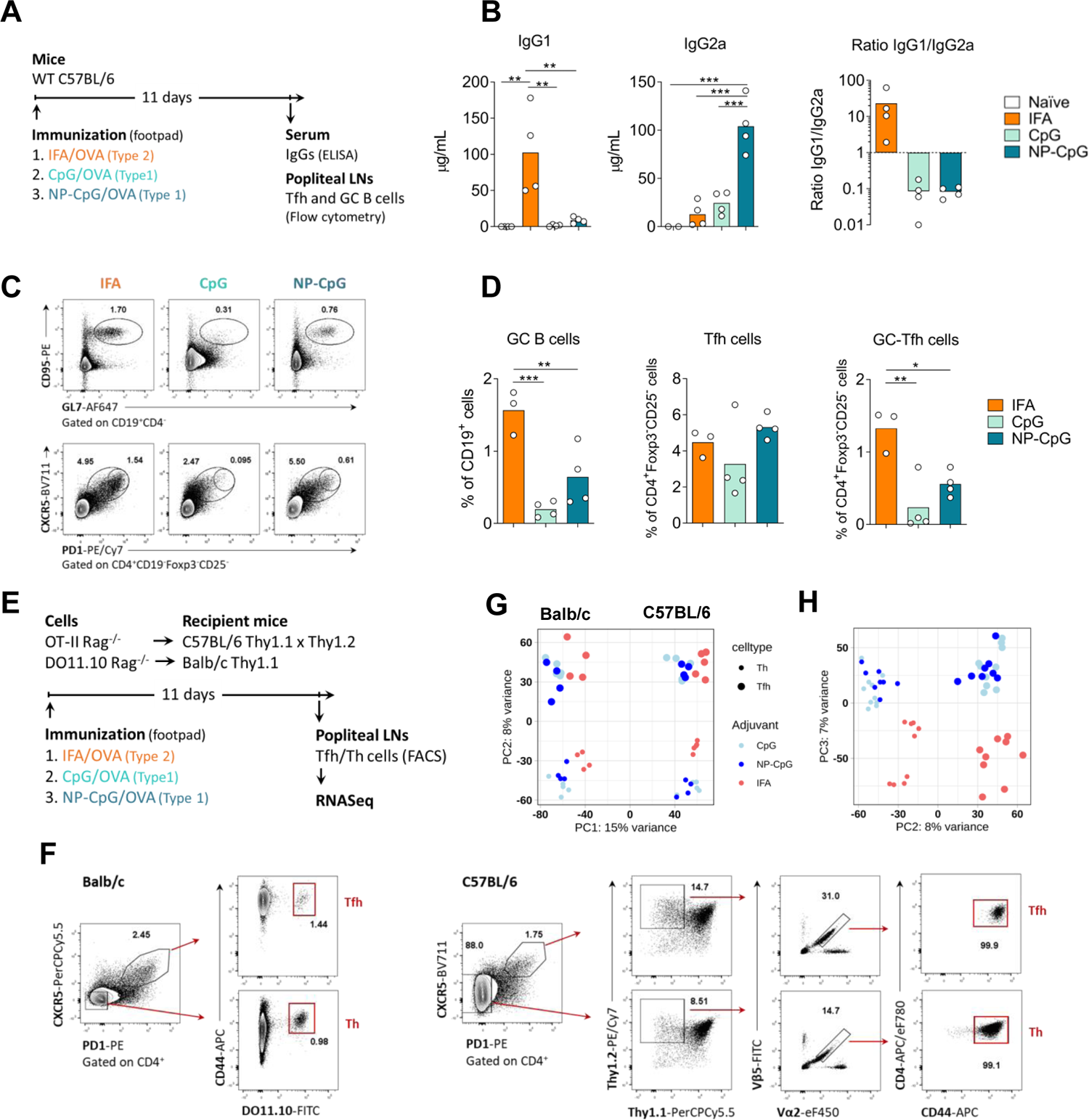
Adjuvants leading to the production of Th1- and Th2-related antibody isotypes induce the differentiation of Tfh cells with distinct transcriptomes. (A) C57BL/6 mice were immunized subcutaneously in the footpad with OVA either emulsified in IFA (IFA), admixed with CpG (CpG), or incorporated with CpG in nanoparticles (NP-CpG). Eleven days later, blood and draining lymph nodes (LN) were collected for analysis. (B) ELISA quantification of anti-OVA IgG1 and IgG2a, and IgG1/IgG2a ratio in serum of the immunized mice. (C) Representative flow cytometry plots and (D) quantification of GC B cells (CD19^+^CD95^+^GL7^+^), Tfh cells (CD4^+^Foxp3^-^CD25^-^CXCR5^+^PD1^+^), and GC-Tfh cells (CD4^+^Foxp3^-^ CD25^-^CXCR5^hi^PD1^hi^) in the draining LNs. Data from one experiment (n = 4), each dot representing one sample and bars representing mean values, analyzed by one-way ANOVA and Tukey’s multiple comparisons tests: **p < 0.01, ***p < 0.001. (E) T cells from OVA-specific TCR-transgenic mice were adoptively transferred into congenic recipients, immunized on the following day with OVA associated with different adjuvants. (F) On day 11, OVA-specific activated Th and Tfh cells were isolated by FACS for RNAseq according to the represented gating strategy. (G) Principal component analysis (PCA) from the RNAseq datasets of Th and Tfh cells, isolated from the two strains, and three types of immunization. PC1 explained 15% of the variance, discriminating datasets from the two strains. (H) PC2 and PC3 have a similar impact on the variance, with PC2 segregating Tfh cells from activated non-follicular T cells, and PC3 separating the samples based on the type of adjuvant used (type-1 vs. type-2).

Immunization with OVA-IFA led to prominent GC responses in the draining lymph nodes (LN), with induction of GC B cells (CD19^+^CD95^+^GL7^+^), Tfh (CD4^+^Foxp3^-^CD25^-^CXCR5^+^PD1^+^), and GC-Tfh (CD4^+^Foxp3^-^ CD25^-^CXCR5^hi^PD1^hi^) cells (**Fig. 1C,D**). Although CpG can directly stimulate B cells, we found that the use of CpG as an adjuvant also led to GC formation and to the emergence of Tfh cells in draining LNs, especially when NP-CpG were used (**Fig. 1C,D**).

To investigate the transcriptome of putative Tfh cells induced under type-1 and type-2 conditions (putative Tfh1 and Tfh2 cells), we used TCR-transgenic T cells to reduce possible sources of variability. In addition, we used two different mouse strains, BALB/c and C57BL/6, known to be biased towards type-2 and type-1 responses, respectively, with the reasoning that the defining characteristics of Tfh1 and Tfh2 cells should be conserved irrespective of the genetic background of the mouse strain. With this approach, we aimed for a near homogenous population of Tfh cells specific for a model antigen (OVA), placing the OVA-specific TCR-transgenic cells under the genetic background of two different mouse strains (**Fig. 1E**). We attempted to preserve the normal physiology by adoptively transferring the TCR-transgenic cells into wild-type congenic hosts before immunization (**Fig. 1E**).

At the peak of the GC response (day 11), we FACS sorted the OVA-specific TCR-transgenic cells from popliteal LNs draining the immunization site. In this way, we obtained near homogeneous populations of OVA-specific Tfh cells (CXCR5^+^PD-1^+^) and activated non-follicular T cells (i.e., CXCR5^-^CD44^+^, referred as Th in the figures) (**Fig. 1F**). Note that in the absence of immunization, the popliteal nodes are devoid of Tfh cells, supporting the idea that virtually all analyzed Tfh cells resulted from the immunization (**Fig. S1**). We sequenced the transcriptome of the OVA-specific Tfh and activated non-follicular T cells from the two strains under the three immunization conditions. In our attempt to sort near-homogenous populations from each immunization, we obtained a small number of cells from each mouse, even at the peak of the GC response. Therefore, we used low input RNA library preparation methods to capture the transcriptome of these samples (see Methods). We generated RNA-seq libraries from 54 samples and sequenced an average of 40 million reads per sample to maximize the capture of the transcriptome of these cells.

Following read mapping and quantification of gene expression, we first performed principal component analysis (PCA) to assess the relationship between the transcriptomes of all the different cell populations (**Fig. 1G, H**). We found that samples sorted from BALB/c and C57BL/6 mice were discriminated by the first principal component, which explains most of the variance (15%), highlighting the strain differences. PC2, with a variance of 8%, discriminated Tfh from non-follicular T cells, and PC3, with a similar variance (7%), described the type-specific segregation of samples (i.e., type-1 versus type-2) (**Fig. 1H**). This segregation of samples shows that a transcriptomic approach can discriminate the cell subsets induced under different types of immunization.

### Transcriptional differences between follicular and non-follicular T cells

Given the clear segregation of Tfh and non-follicular T cells in both strains, as observed in the PCA (**Fig. 1H**), we investigated the transcriptional differences between the two cell subsets in both mouse strains. Differential gene expression analysis revealed 702 significantly differentially expressed (DE) genes. *Cxcr5, Pdcd1, Il21, Bcl6, Il1r1,* and *Sh2d1a* were upregulated in Tfh cells compared to activated non-follicular T cells. These genes were described as hallmarks of the Tfh phenotype (*20, 24*). In contrast, *Ccr7, S1pr1, Sell, Klf2, and Selplg* – genes that restrict GC entry and are inhibitory for the Tfh phenotype – were upregulated in non-follicular T cells (**Fig. 2A**). We obtained a list of genes annotated as cytokines, chemokines, interferons, interleukins, and their receptors, including TNF and TGF beta family members, from the ImmPort database (*25*) (referred from now on as the Immune gene list), and compared this list against the significantly differentially expressed genes (DEGs). We found the expression of well-described cytokines and chemokines in line with the known differences between Tfh and non-follicular T cells (**Fig. 2B**).

**Figure 2.**
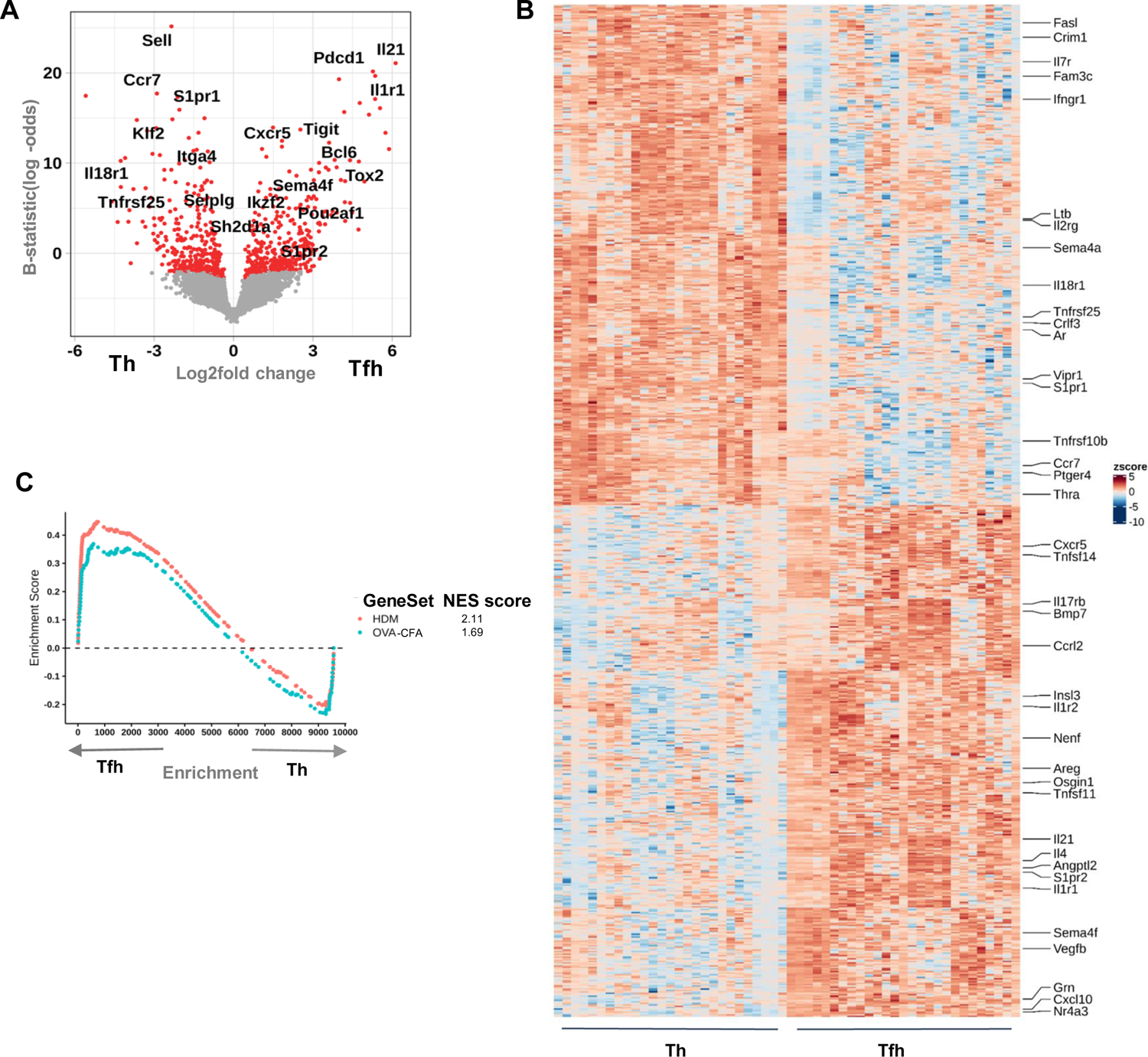
Tfh and activated non-follicular T cells have a distinct transcriptome. **(A)** Volcano plot of all significantly differentially expressed genes between Tfh and activated non-follicular T cells. A selection of Tfh-specific and non-follicular T cell-specific genes are labeled. Genes with adjusted p values of less than 0.05 were considered significant and are represented in red. **(B)** Heatmap of the most significantly differentially expressed genes between the two cell populations, with the genes matching the ImmPort immune gene list labeled. **(C)** Gene set enrichment analysis of our Tfh datasets against publicly available datasets of Tfh samples (GSE134153). The Normalized Enrichment Score (NES) of both samples with a significant adjusted p-value of less than 0.05 is shown.

We performed a gene set enrichment analysis (GSEA) of our samples against publicly available datasets of Tfh samples generated in mice exposed to OVA-complete Freund’s adjuvant (CFA) immunization or allergic disease induced with House Dust Mite (HDM) (*26*). The Tfh transcriptome from our data largely matched the Tfh transcriptome from the two public datasets, further confirming the phenotype of our samples as *bona fide* Tfh cells (**Fig. 2C**). Since the PC variance of 8% between Tfh and non-follicular T cell subsets was able to correctly re-capitulate the Tfh transcriptome, we concluded that these datasets were adequate to explore the characteristics of putative Tfh1 and Tfh2 subsets.

### Transcriptional Signatures of Tfh1 versus Tfh2 cells

We next performed differential gene expression analysis of Tfh samples generated from type-1 and type-2 immunizations. We analyzed 8888 genes, of which 467 resulted as significantly differentially expressed genes (**Fig. 3A**). Among those were genes implicated in type-2 responses (namely *Il4* or *Cebpb*) upregulated in Tfh2 samples, while *Sema4a* was upregulated in Tfh1 samples (*27, 28*). We then compared our ImmPort immune gene list against the Tfh1 vs. Tfh2 differentially expressed genes (**Fig. 3B and Fig. S2A**). It should be noted that while certain genes (e.g*. Il21, Il1r1, Tnfsf11*) appear to be expressed only in Tfh2 samples (**Fig. 3B**), the unscaled plot indicates that these molecules are expressed in all Tfh samples, but at different intensities between the two subsets (**Fig. S2A**).

**Figure 3.**
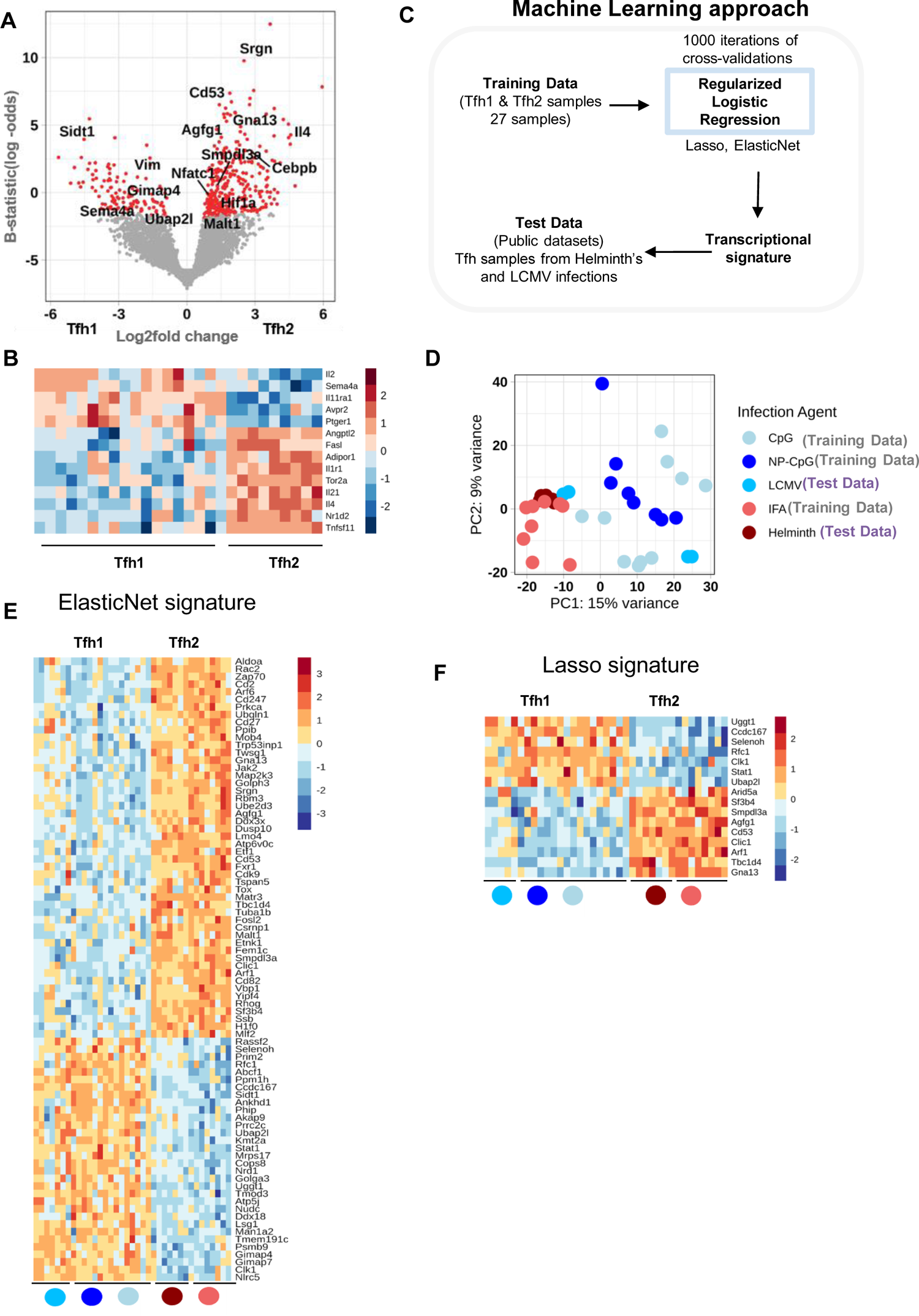
The transcriptional signature of Tfh1 and Tfh2 subsets. **(A)** Volcano plot of all significantly differentially expressed genes between Tfh1 and Tfh2 samples. Genes with adjusted p values of less than 0.05 were considered significant and are represented in red. **(B)** Heatmap of genes matching the ImmPort immune gene list and significantly differentially expressed between the two subpopulations. **(C)** The workflow design of the machine learning approach used to generate the transcriptional signatures of the Tfh1 and Tfh2 datasets. **(D)** Principal Component Analysis of all training and test datasets (GSE105808, GSE79039, GSE72568) highlighting the heterogeneity of all datasets. **(E)** Heatmap of the 82 signature genes identified using ElasticNet penalty. The heatmap shows a similar profile in the training and test datasets for Tfh1 and Tfh2 populations. **(F)** Heatmap of the more concise signature of 16 genes identified using Lasso penalty, that can accurately classify training and test datasets as Tfh1 and Tfh2. The Lasso signature is mostly a subset of the ElasticNet signature. The heatmap also shows the consistency of the signature between training and test datasets. The colored dots in **(E-F)** refers to the test and training datasets as in **D**.

Although we saw a preferential expression of some transcripts associated with the different types of responses (namely, *Il4* on putative Tfh2; and *Sema4a* on putative Tfh1, **Fig. 3B**), none of the transcripts could uniquely discriminate between the Tfh populations induced under the different responses. In addition, differential gene expression analysis did not yield any single gene that could uniquely identify Tfh cells as type-1 or type-2. Instead, a collection of genes showed overall differential expression patterns (**Fig. S2B**). Therefore, instead of a single-gene definition for Tfh1 or Tfh2 cell subsets, a transcriptional signature of a collection of genes appears to be a better approach to defining these populations.

We used a machine learning approach to define a minimal signature for Tfh1 and Tfh2 cell subsets (**Fig. 3C**). We trained a logistic regression model with two different regularization penalties (ElasticNet and Lasso) to generate a transcriptional signature for Tfh1 and Tfh2 cell subsets from our datasets. While the resulting signatures could classify our samples correctly, this could be due to overfitting and required further validation, ideally with independent datasets of Tfh1 and Tfh2. Thus, to test the signature’s robustness independently from our training datasets, we collected publicly available transcriptome datasets of Tfh samples generated through murine infection with helminths (a type-2 infection) or LCMV (a type-1 infection). These datasets were generated in different laboratories, obtained on different days post-infection, independently from the TCR-transgenic cells we used, and collected from a distinct lymphoid tissue (the spleen). A PCA visualization of all training and test data samples highlights their heterogeneity (batch corrected) (**Fig. 3D**). We found that the transcriptomic signatures generated using our samples (mice immunized with NP-CpG, CpG, and IFA) were accurate enough to correctly classify all public datasets as either type-1 or type-2 (**Table S3**). ElasticNet led to the identification of a transcriptional signature of 82 genes, while Lasso restricted the signature to 16 genes. Furthermore, a heatmap of the transcriptional signature genes in both training and test datasets showed highly similar patterns of expression between the training datasets (with adjuvant) and the test datasets (with infection) (**Fig. 3E-F**). Finally, we examined the correlation between the differentially expressed gene analysis of Tfh1 and Tfh2 samples against the transcriptional signature, and we found a clear correlation among them (**Fig. S2C**). These results show that Tfh1 and Tfh2 cells comprise two distinct Tfh populations characterized by different transcriptional programs.

### Functional specialization of Tfh cells upon type-1 and type-2 immunization

We next investigated the functional outcome of Tfh1 and Tfh2 cells, sorting these populations from mice immunized under type-1 and −2 conditions as described above. Given the low number of Tfh1 and Tfh2 cells in popliteal LN, we pooled cells from different animals. We co-cultured sorted Tfh cells and B cells, stimulated with anti-CD3 and anti-IgM (**Fig. 4A**). At the end of the culture, we found that supernatants from cultures with Tfh cells isolated from mice immunized with OVA/IFA (type-2) contained more IL-4, while cultures with Tfh cells derived from OVA/NP-CpG (type-1) immunization produced more IFNγ (**Fig. 4B**).

**Figure 4.**
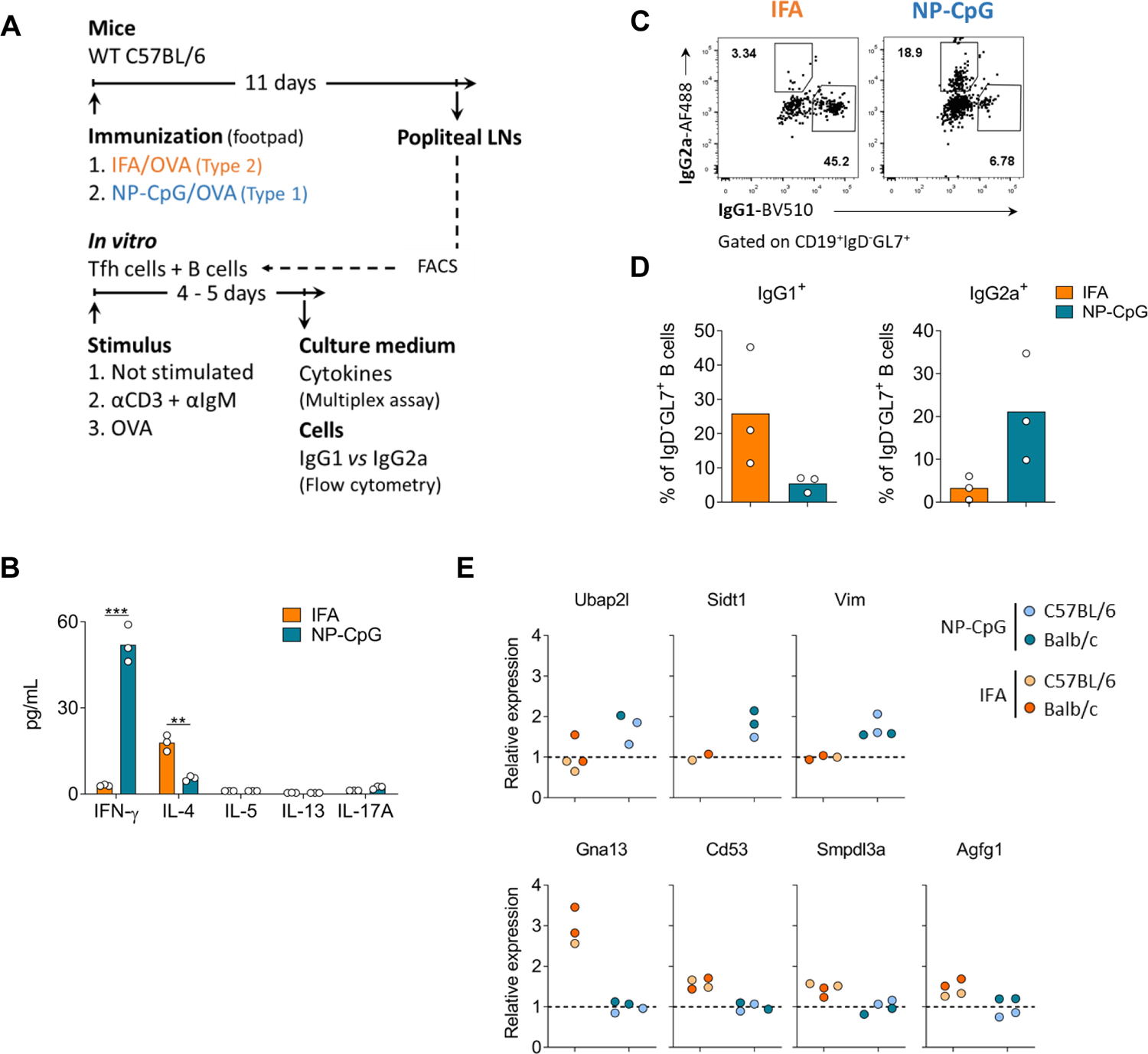
Tfh1 and Tfh2 cells are functionally distinct. **(A)** C57BL/6 mice were immunized in the footpad with OVA emulsified in IFA (IFA) or incorporated with CpG in nanoparticles (NP-CpG). On day 11, Tfh (CD4^+^CD25^-^CXCR5^+^PD1^+^) and B cells (CD19^+^CD4^-^) were isolated from draining LNs by flow cytometry and co-cultured. **(B)** After 5 days of culture, cytokines in the culture medium were quantified by multiplex assays. Data representative from two experiments (culture triplicates performed with cells obtained from 10 immunized mice per group), each dot representing one replicate and bars representing mean values, analyzed by Student’s *t*-test: **p < 0.01, ***p < 0.001. **(C)** Representative dotplots and **(D)** quantification of IgG2a^+^ and IgG1^+^ isotype-switched B cells at the end of the co-cultures (stimulation with OVA), analyzed by flow cytometry. Data from one experiment (culture triplicates performed with cells obtained from 10 immunized mice per group), each dot representing one replicate and bars representing mean values, analyzed by Student’s *t*-test: **p < 0.01, ***p < 0.001. **(E)** Evaluation of gene expression by quantitative real-time PCR in Tfh cells isolated from mice 11 days after immunization under type-1 and type-2 conditions as described in (A). The cells from 10 mice immunized with each adjuvant were pooled, and the resulting cDNA was tested in duplicate, both from C57BL/6 and Balb/c mouse strains. 2^-ΔCT^ values were determined in reference to the *Actb* housekeeping gene of the same sample and then normalized to the average 2^-ΔCT^ values obtained for mice immunized with the other adjuvant.

Furthermore, in the cultures with Tfh cells from mice immunized with IFA (putative Tfh2 cells) and stimulated with OVA, the B cells displayed a preferential isotype switching towards IgG1. In contrast, cultures with Tfh cells from NP-CpG immunized mice favored isotype switching towards IgG2a (**Fig. 4C,D**).

The functional polarization of Tfh cells isolated from LN draining the immunization site was aligned with the anticipated phenotypic changes that emerged from the transcriptional signature deduced above. We found that Tfh cells isolated from mice immunized under type-1 conditions had more significant expression of *Gimap4, Ubap2l, Sidt1,* and *Vim*, while conversely, Tfh cells arising from type-2 immunizations displayed greater expression of *Malt1, Gna13, Cd53, Smpdl3a*, and *Agfg1* (**Fig. 4E**). These changes were consistent in BALB/c and C57BL/6 genetic backgrounds (**Fig. 4E**).

### Single-cell transcriptomes of Tfh cells induced under type-1 and type-2 immunization

The results described above defined clear Tfh1 and Tfh2 transcriptional signatures. However, immunization with type-1 and −2 adjuvants leads to the predominant production of immunoglobulins of the selected type and a small amount of immunoglobulin of the divergent type (**Fig. 1B**). This observation led us to investigate the hypothesis of the presence of a small proportion of Tfh cells with divergent functional specialization. We had to rely on a method able to identify the characteristics of individual cells to address this issue. We generated single-cell RNA-seq datasets from Tfh cells sorted from Foxp3*^gfp^* reporter mice under NP-CpG and IFA immunizations (**Fig. 5A**). While in previous experiments with adoptively transferred TCR-transgenic T cells, virtually all Tfh cells were devoid of Foxp3 (i.e., without Tfr cells), there is a significant number of Tfr cells in a wild-type population. To exclude the Tfr cells from the analysis of Tfh subsets, we enriched for either CXCR5^+^Foxp3*^GFP^*^-^ or CXCR5^-^ Foxp3*^GFP^*^+^ cells (**Fig. S3A**). The sequenced Treg cells facilitate the identification of *bona fide* Tfr cells, as we have shown in a recent study with human cells (*29*). The assessment of immunoglobulin production confirmed the different types of response: higher IgG1 in OVA/IFA immunization (type-2 response) and predominant IgG2a production with OVA/NP-CpG (type-1 response) (**Fig. S3B**).

**Figure 5.**
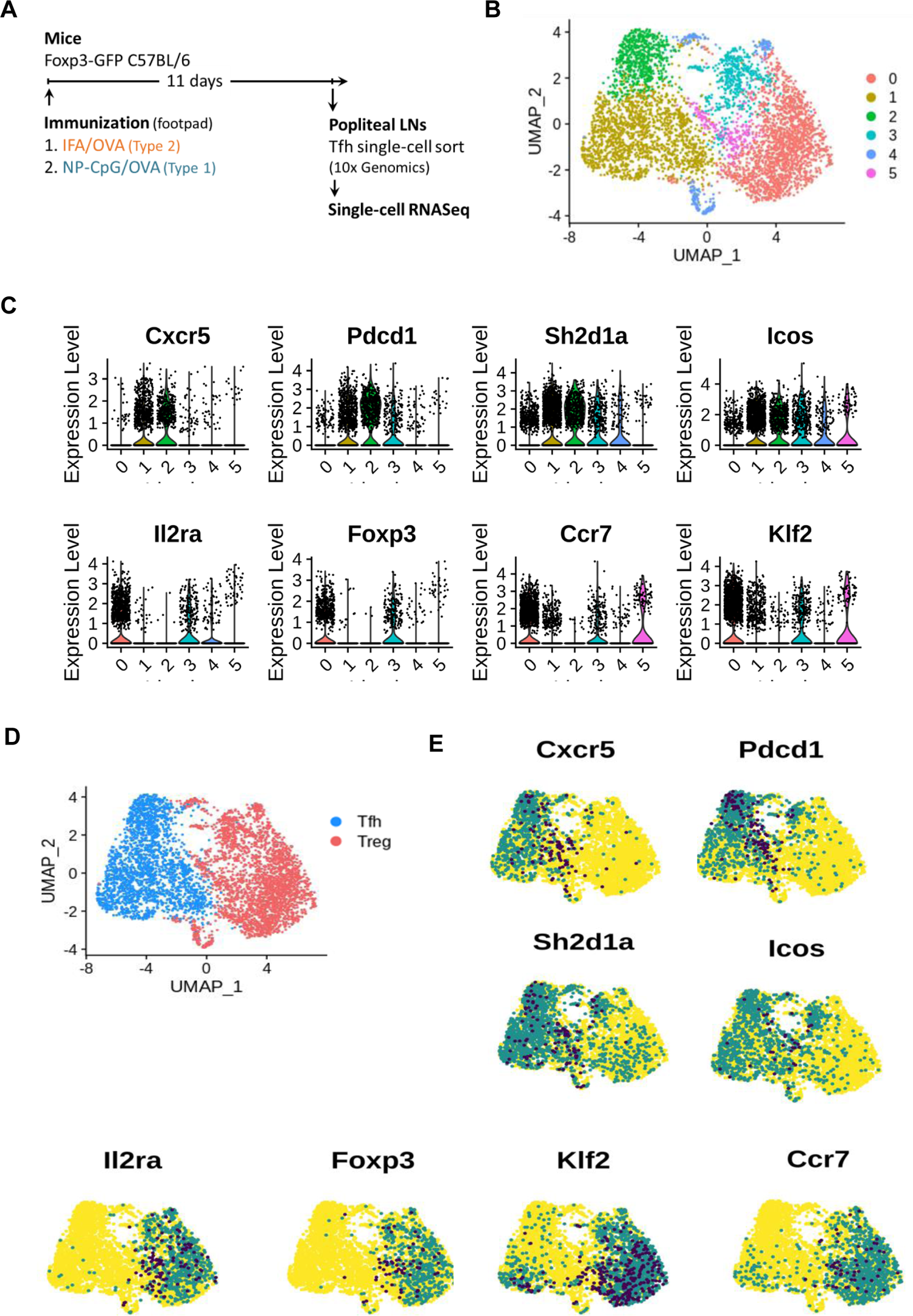
The single-cell transcriptome of Tfh cells. **(A)** C57BL/6 Foxp3*^GFP^* reporter mice were immunized subcutaneously in the footpad with OVA in IFA or NP-CpG. Eleven days later, draining lymph nodes were collected for analysis. T cells were sorted as represented in Fig. S2. **(B)** UMAP projection showing the clustering of all sorted T cells in six clusters. **(C)** Violin Plots of the expression level of transcripts associated with Tfh and Treg cells in each cluster. **(D)** UMAP projection of all cells categorized as belonging to either Tfh or Treg clusters, according to their transcriptional profile. **(E)** Feature plots of known genes associated with Tfh and Treg populations, showing a consistent expression for the respective cell populations, confirming the identity of the clusters.

After quality control, the transcriptome from 4918 single-cells was analyzed, which resulted in six main clusters (**Fig. 5B**). The expression of the hallmark Tfh genes *Cxcr5, Pdcd1, Sh2d1a* (encoding SAP), and *Icos* was highest in clusters 1 and 2, while expression of the Treg-associated genes *Foxp3, Il2ra* (encoding CD25), *Ccr7*, *and Klf2* was observed in clusters 0, 3, 4, and 5 (**Fig. 5C**). Using additional Treg and Tfh markers, we confirmed the identity of cells from clusters 0, 3, 4, and 5 as Treg cells and clusters 1 and 2 as Tfh cells (**Fig. S4A-B**). We, therefore, renamed these clusters as belonging to either Tfh or Treg categories (**Fig 5D-E**).

### Heterogeneity of Tfh cell populations generated under type-1 and type-2 immunizations

We next analyzed the combined Tfh cells from the two immunizations, excluding the Foxp3^+^ cells. We performed an unbiased low-resolution clustering to capture the global profile of these cells and found three main clusters (**Fig. 6A**). To identify the distinctive markers for each cluster, we assessed the expression of genes listed in the immune gene list. We found cluster 0 showed high expression of *Cxcr3* and *Ifng*, while cluster 1 showed increased expression of *Il4* (**Fig. 6B**), confirming the identity of cluster 0 as Tfh1, and cluster 1 as Tfh2. We also evaluated the expression of known Tfh transcripts in both clusters. We found a consistent expression of Tfh-related transcription factors in both Tfh1 and Tfh2 clusters, confirming the common follicular profile of these cells (**Fig. S5**). Cluster 2 cells showed expression of cell cycle markers, namely *Mki67* and *Top2a*.

**Figure 6.**
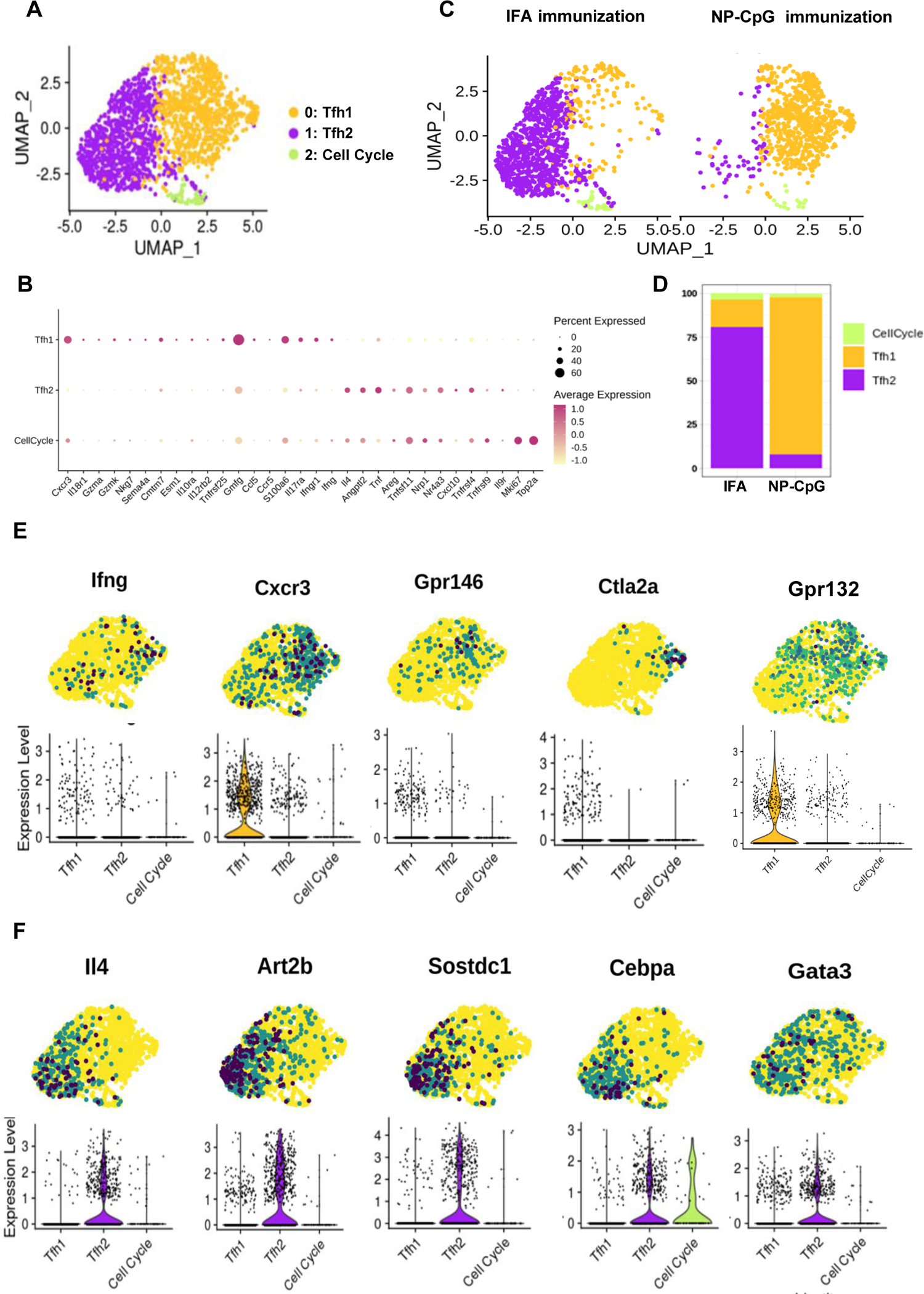
Heterogeneity of Tfh cells under the two types of immunization. **(A)** UMAP projection of only the Tfh cells, re-analyzed without Treg cells, showing clustering as Tfh1, Tfh2, and a minor population with high expression of cell cycle genes, labeled as Cell Cycle. **(B)** Expression of the immune gene list for each Tfh sub-population labeled as in (A). **(C)** UMAP projection of the cells defined in (A) plotted based on the type of immunization. The LNs from mice immunized with a type-2 adjuvant (IFA) had a majority of Tfh cells classified as Tfh2 cells (left), while mice immunized with a type-1 adjuvant (NP-CpG) had a majority of Tfh1 cells (right). **(D)** Percentage of Tfh1 and Tfh2 cells found in each immunization. **(E)** Feature plots and violin plots representing the expression level of genes associated with the Tfh1 cells, or **(F)** with the Tfh2 cells. All represented genes were significantly differentially expressed between the Tfh1 and Tfh2 cell subsets.

Importantly, we found the clustering of Tfh cells showed segregation of cells based on the immunizing adjuvant: type-2 immunization (IFA) led to a predominance of Tfh2, and type-1 immunization (NP-CpG) to Tfh1 (**Fig. 6C**). Examining the proportion of the different cell subsets in each immunization, we found that NP-CpG samples (type-1) contained ∼90% of Tfh1 cells with a minor population of ∼8% of Tfh2 cells (**Fig. 6D**). On the contrary, IFA samples (type-2) comprised ∼80% of Tfh2 cells with a smaller population of ∼15% of Tfh1 cells. Both immunizations also showed ∼2-3 % of cells undergoing cell cycle (**Fig. 6D**). Differential gene expression between the Tfh1 and Tfh2 clusters identified genes we had validated from bulk-RNAseq, and also some additional new genes, such as *Cxcr3* for Tfh1 and *Cebpa* for Tfh2 cells, able to discriminate the two populations (**Fig. 6E-F**).

Overall, using single-cell resolution, we were able to dissect the heterogeneous profile observed in the bulk RNA-seq datasets by identifying the proportion of Tfh1 and Tfh2 subsets generated from each immunization. Furthermore, this cell-based approach led to the identification of additional markers with a preference for Tfh1 and Tfh2 cells.

### Germinal centres induced upon type-1 or −2 immunization are enriched in Tfh1 or Tfh2 cells

Finally, we used the additional transcripts identified following scRNAseq to directly visualize Tfh1 and Tfh2 cells within LNs of immunized mice. We further confirmed that Tfh cells sorted from LNs draining the site of immunization with type-1 (NP-CpG) or type-2 (IFA) adjuvants showed a preferential expression of *Ctla2a* and *Cxcr3* (type-1); and *Hif1a, Cebpb, Gata3, Cd200*, and *Nfatc1* (type-2) through RT-qPCR (**Fig. 7A**). Additionally, we used flow cytometry to establish a distribution of CXCR3, CD53, CD200, and LAG3 in Tfh1 and Tfh2 cells consistent with the gene expression (**Fig. 7B**).

**Figure 7.**
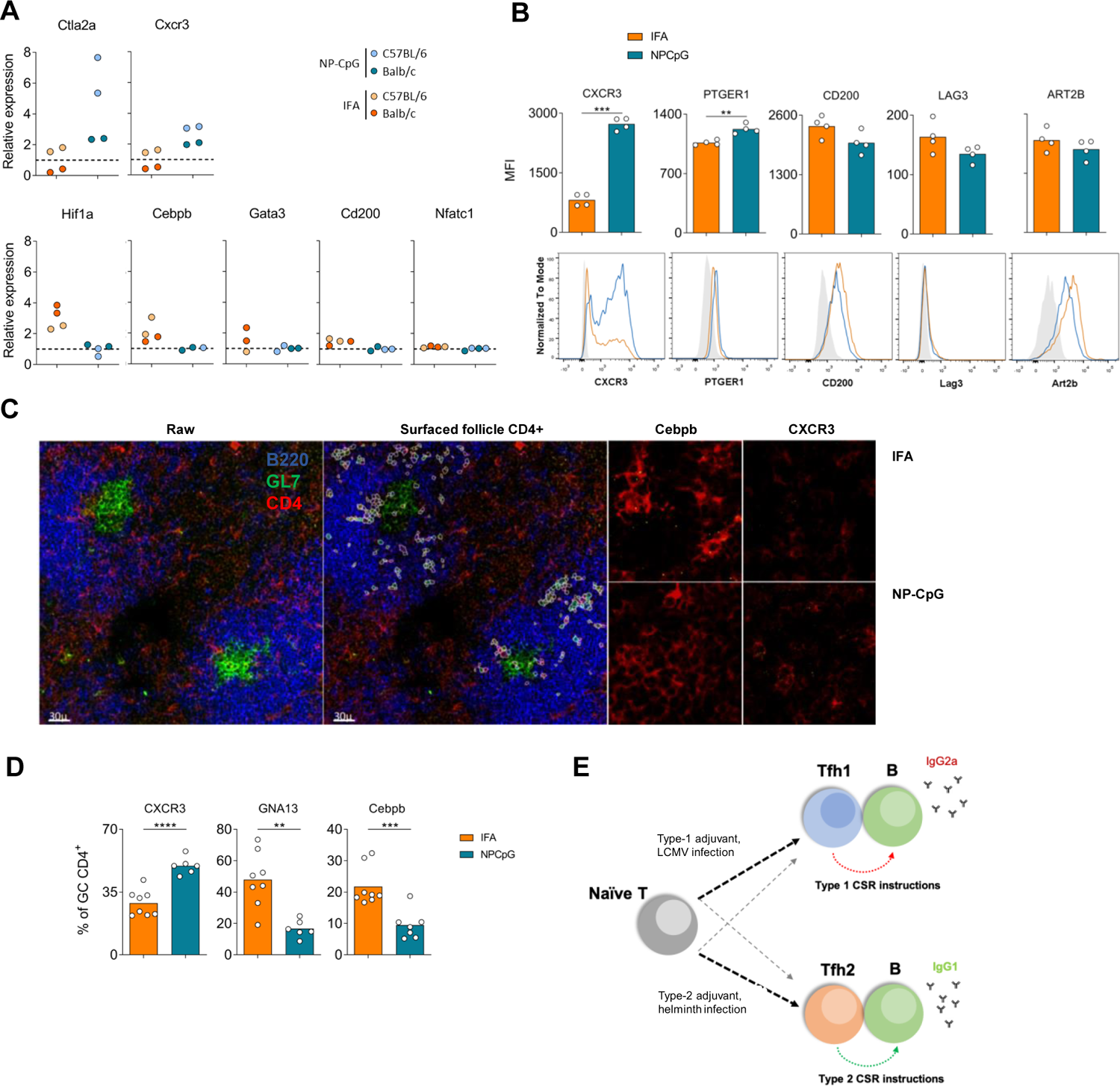
Germinal centres induced upon type-1 or −2 immunization are enriched in Tfh1 or Tfh2 cells. **(A)** Mice were immunized in the footpad with OVA emulsified in IFA (IFA) or incorporated with CpG in nanoparticles (NP-CpG), as described in Fig. 1, and the expression of selected genes by Tfh cells sorted 11 days after immunization was evaluated by quantitative real-time PCR. The cells from 10 mice immunized with each adjuvant were pooled and tested in duplicate, both from C57BL/6 and Balb/c mouse strains. 2^-ΔCT^ values were determined in reference to the *Actb* housekeeping gene of the same sample and then normalized to the average 2^-ΔCT^ values obtained for mice immunized with the other adjuvant. **(B)** The expression of selected genes was further evaluated by flow cytometry in Tfh cells from C57BL/6 mice immunized in a similar manner. Representative histograms and quantification of median fluorescence intensity (MFI) of Tfh cells (CD4^+^Foxp3^-^CD25^-^CXCR5^+^PD1^+^) in the draining lymph nodes from mice immunized with IFA (orange) or NP-CpG (blue). Data from one experiment (n = 4), each dot representing one sample and bars representing mean values, analyzed by Student’s *t*-test: **p < 0.01, ***p < 0.001. **(C)** Representative raw image of stained draining LN of mice, 11 days following immunization with IFA or NP-CpG as described, alongside example segmentation of CD4^+^ cells within B cell follicles. Representative probe staining for Cebpb and CXCR3 are shown for each group. **(D)** The proportion of germinal centre CD4^+^ T cells positive for each marker was quantified for images processed as in (C) and quantified. Data from one experiment (n=3), each dot representing one replicate and bars representing mean values, analyzed by Student’s *t*-test: **p < 0.01, ***p < 0.001, ****p<0.0001. **(E)** Model representing the specialization of Tfh cells under type-1 or −2 conditions. Type-1 adjuvants or LCMV infection drives specialization of the majority of Tfh cells towards Tfh1, and a minority of Tfh2 cells. This Tfh1 cell specialization favors IgG2a isotype switching. Conversely, type-2 adjuvants or Helminths’ infection drive mostly Tfh2 specialization, with a small percentage of Tfh1 cells and a predominant production of IgG1.

Then, we used RNAscope to investigate the characteristics of GC Tfh cells within the draining LNs from mice immunized with the two types of adjuvants. This strategy allowed us to find that LNs from mice immunized with a type-1 adjuvant (NP-CpG) were preferentially harboring Tfh cells displaying *Cxcr3* transcripts, whereas type-2 immunization (IFA) led to a preferential accumulation of Tfh cells within the GCs containing *Gna13* and *Cebpb* transcripts (**Fig. 7C,D**). The direct visualization of transcripts in Tfh cells from popliteal LNs draining the immunization site firmly demonstrated that type-1 and −2 adjuvants drive the preferential participation of Tfh1 and Tfh2 cells, respectively, in humoral responses.

## Discussion

In this work, we aimed to investigate how a putative functional specialization of Tfh cells could explain the selection of appropriate humoral responses classically attributed to Th1 and Th2 subsets. The study of Th1 and Th2 polarization was greatly facilitated by *in vitro* assays leading to the functional polarization of the two subsets under very controlled conditions (*18*). By contrast, a major difficulty in studying Tfh cells under type-1 or −2 conditions has been the lack of appropriate *in vitro* assays for Tfh cell differentiation. To overcome this difficulty, we created a controlled *in vivo* experimental system, using an adjuvant-based immunization strategy, to generate comparable Tfh cells biased towards either type-1 or type-2 responses.

We immunized mice in the footpad and collected Tfh cells generated from adoptively transferred TCR transgenic cells in popliteal LN to maximize the homogeneity in the type-specificity of Tfh cell populations. The popliteal nodes from non-immunized mice do not have Tfh cells, and the adjuvants (IFA and CpG) are devoid of additional proteins. As a result, we could be certain that the Tfh cells (and control TCR-transgenic non-Tfh cells) were induced in response to the immunization with the distinct adjuvants. In addition, it is established that C57BL/6 mice are more prone to type-1 responses, while BALB/c mice favor type-2. To avoid capturing strain-biased responses of type-1 and type-2 Tfh cells, we used those two strains of mice with distinct preferences for type-1 and −2 responses. The drawback of the homogeneity of the Tfh cells was the very low number of cells to analyze, requiring sequencing methods appropriate for low cell yield.

The *in vivo* strategy to obtain the transcriptome of homogeneous populations of Tfh cells, coupled with a machine learning approach, established the transcriptional signature of Tfh1 and Tfh2 cells. We found this signature consistent with publicly available datasets of Tfh samples from different laboratories generated using infection models.

The notion of different subpopulations of Tfh cells, namely Tfh1 and Tfh2, has already been investigated (*30*). The first reports on functional Tfh subsets relied on CXCR3 and CCR6 to define human Tfh1 (CXCR3^+^CCR6^-^), Tfh17 (CXCR3^-^CCR6^+^), and Tfh2 (CXCR3^-^CCR6^-^) in the blood (*31*), but the same markers failed to identify equivalent Tfh subsets within human lymphoid tissue (*30, 32*). Furthermore, blood Tfh1 cells, defined as CXCR3^+^CCR6^-^, lacked effective helper function and were suggested to have a suppressive role (*30, 31, 33–35*). Other studies relied on the expression of IFNγ or IL-4 (namely, using an IL-4 reporter system in mice) to identify the Tfh cells associated with type-1 or −2 conditions (*12–14*). Nevertheless, the two subsets – Tfh1 and Tfh2 – were never addressed comparatively in those studies. A different functional subset of Tfh cells, the Tfh13, was described in animal models of allergic disease (*36*). Therefore, our study provides a much-needed direct comparison of Tfh cells induced in the tissue under type-1 and −2 conditions.

It should be noted that our results show that a single gene approach does not accurately distinguish Tfh1 from Tfh2 cells. The machine learning approach provided the necessary gene signature for accurately classifying a given sample as Tfh1 or Tfh2. But the classifiers use a minimal number of genes that can accurately classify the samples and do not represent the complete transcriptional profile of the two subsets. For example, if several genes are coordinately expressed, the machine learning algorithm will only retain one of the genes for the signature as the additional genes do not provide added information for the accurate classification of the cells. It is, therefore, unsurprising that our subsequent scRNAseq studies showed additional genes differentially expressed between Tfh1 and Tfh2 cells.

The analysis of the bulk populations did not address a possible degree of heterogeneity in the two immunizations, as this can only be studied at a single-cell level. To investigate this, we generated single-cell transcriptomics datasets under similar immunization conditions. We found ∼10-15% of Tfh cells divergent from the immunization type (i.e. a small proportion of Tfh2 cells in mice subjected to type-1 immunization and vice-versa). This finding is in accordance with the antibody titers observed in the two immunizations, where it is common to find a small proportion of antibodies of the divergent type.

We observed that Tfh cells isolated from mice immunized under type-1 and −2 conditions displayed a distinct functional behavior confirmed through *in vitro* assays, leading to the provision of appropriate help to B cells biased to the appropriate type. Furthermore, the direct visualization of LN allowed the observation of Tfh cells within the GC expressing transcripts associated with their functional specialization in Tfh1 and Tfh2. We did not directly assess whether polarized Tfh cells differentiate from polarized Th cells or, alternatively, from naïve CD4 T cells. However, there is evidence that Tfh and Th1 cells follow a bifurcated differentiation trajectory from naïve CD4 T cells (*37*). Therefore, it is likely that the emergence of specialized Tfh1 and Tfh2 cells occurs in parallel with Th1 and Th2 polarization. It has been described that the strength of TCR ligation or IL-2 availability can drive the decision between Tfh versus Th commitment (*38–41*). It is possible that Th1 and Th2 polarization is not completely dissociated from Tfh1 and Tfh2. In the same way that cytokines from Th1 reinforce T cell polarization to Th1 fate, and Th2 cytokines to Th2 fate, it is conceivable that a similar impact of Th1 and Th2 cytokines can favor the differentiation towards, respectively, a Tfh1 and Tfh2 fate.

Our results support a model for Tfh specialization according to the existing polarizing conditions (**Fig. 7E**), where type-1 and type-2 immunization leads to the emergence of specialized Tfh1 and Tfh2 cells of the concordant type, along with a minor proportion of Tfh cells of the divergent type.

In summary, our results provide compelling evidence for the functional specialization of Tfh1 and Tfh2 subsets under type-1 or −2 immune responses. The definition of the transcriptional profile of Tfh1 and Tfh2 cells offers new targets for therapeutic modulation of GC responses targeting specifically type-1 or type-2 humoral immunity.

### Methods and Materials

#### Mice and animal procedures

The experimental plan relied on mice from the following strains: C57BL/6, C57BL/6 Thy1.1 x Thy1.2, OT-II.Rag-/-, Balb/c, Balb/c Thy1.1, DO11.10.Rag-/-, and C57BL/6 Foxp3*^GFP^* reporter mice. The mice were bred, maintained under specific pathogen-free conditions, and used at the iMM under an animal experimentation authorization granted by the ORBEA-iMM (iMM’s Animal Welfare Body) and DGAV (the Portuguese National Authority for Animal Health) and followed European Union guidelines.

Mice aged between 8 to 12 weeks were immunized subcutaneously in the footpad with ovalbumin (Ovalbumin EndoFit, Invivogen, #vac-pova) either emulsified 1:1 (v:v) with IFA (IFA, Sigma-Aldrich, #F5506), admixed with CpG (ODN 1826 - TLR9 ligand, Invivogen, #tlrl-1826), or entrapped with CpG in polymeric nanoparticles (*42*). Each animal was inoculated in the paw with one of the antigen/adjuvant mixtures, with a volume of 50 μL per paw containing 80 μg of OVA and, in the case of CpG or NP-CpG formulations, 30 μg of CpG. Popliteal lymph nodes and blood were collected on day 11 following immunization.

#### Cell sorting and flow cytometry analysis

Single-cell suspensions were obtained by disrupting the lymph nodes in PBS (Lonza) with 2% FBS (Gibco) with curved forceps and a nylon mesh. Surface stainings were performed in the same buffer with the following monoclonal antibodies or reagents: anti-Vβ5.1,5.2 TCR (MR9-4, BD Bioscience), anti-CD279 (PD-1) (J43, eBiosciences), anti-Thy1.1 (HIS51, eBiosciences), anti-Thy1.2 (53-2.1, eBioscience), anti-CD44 (IM7, Biolegend), anti-CD4 (RM4-5, eBioscience), anti-CD4 (RM4-4, Biolegend), CD4 (GK1.5, eBioscience), anti-Vα2 TCR (B20.1, eBioscience), anti-CD185/CXCR5 (2G8, BD Bioscience), anti-TCR DO11.10 (KJ1-26, eBiosciences), anti-CD19 (MB19-1, eBioscience), anti-CD8 (53-6.7, eBioscience), anti-CD25 (PC61.5, eBioscience), anti-CD19 (1D3, eBioscience), anti-I-A/I-E (M5/114.15.2, Biolegend), anti-IgD (11-26c.2a, BioLegend), GL7 (eBioscience), anti-CXCR3 (CXCR3-173, BioLegend), anti-PTGER1 (#orb103299, Biorbyt), anti-CD200 (OX90, eBioscience), anti-ART2B (Nika102, Novus Biologicals), anti-LAG3 (C9B7W, eBioscience), and Streptavidin (Biolegend). The LIVE/DEAD Fixable Aqua Dead and Near-IR Cell Stain Kits (Molecular probes, Life Technologies) or DAPI (Biolegend) were used for dead-cell exclusion. For some flow cytometry analysis, in addition to the surface staining with the mentioned antibodies, intracellular staining was performed after fixation and permeabilization with the Foxp3/Transcription Factor Staining Buffer Set Foxp3 Staining Set (eBioscience, #00-5523-00), according to the manufacturer’s instructions. Antibodies used for intracellular staining were: anti-Foxp3 (FJK-16s, eBioscience), IgG2a-AF488 (RMG2a-62, Biolegend), and IgG1-BV510 (RMG1-1). Cells were sorted on a BD FACSAria cell sorter and flow cytometry analysis was done on a BD LSR Fortessa flow cytometer. Acquisition data were analysed on FlowJo software (Tree Star).

#### ELISA for immunoglobulin quantification

OVA-specific immunoglobulin concentration in the serum was determined by ELISA. Briefly, high protein-binding ELISA plates (Nunc MaxiSorp, #44-2404-21) were coated overnight at 4°C with OVA (Invivogen, #vac-pova) at 10 µg/ml in coating buffer (eBioscience, #00-0044-59). Serum samples and mouse anti-OVA immunoglobulins used to generate standard curves (anti-OVA IgG1, clone L71, #7093; anti-OVA IgG2a, clone M12E4D5, #7095; and anti-OVA IgG2c, clone 3E3A9, #7109; all from Chondrex) were serially diluted in assay buffer (eBioscience ELISA/ELISPOT Diluent, #00-4202-56) and incubated overnight at 4°C. The plates were then incubated 1 hour at room temperature with goat anti-mouse immunoglobulins conjugated with horseradish peroxidase (HRP) as detection antibodies (anti-IgG1, #1070-05; anti-IgG2a, #1080-05; anti-IgG2c, #1077-05; all from SouthernBiotech). Finally, the chromogenic HRP substrate 3,3’,5,5’-tetramethylbenzidine (TMB, eBioscience, #00-4201-56) was added and the reaction was stopped with H_2_SO_4_. The color development was measured through spectrophotometric absorbance at 450 nm. The dilutions of serum samples showing optical densities falling within the standard curve values were used for quantification.

#### Quantitative PCR

C57BL/6 and Balb/c mice were immunized in the footpad with OVA in IFA or in NP-CpG and single cell suspensions were prepared 11 days later from popliteal lymph nodes pooled from 8 to 10 mice per group. Total RNA was purified from FACS-sorted Tfh cells (CD4^+^CD25^-^CXCR5^+^PD1^+^CD44^+^) using the RNeasy Plus Micro Kit (Qiagen, #74034) and cDNA was synthesized with the SuperScript IV First-Strand cDNA Synthesis Reaction kit (Invitrogen, #18091050) according to manufacturer’s instructions. Each sample was tested in duplicate for the expression of selected genes in the ViiA 7 real-Time thermal cycler (Applied Biosystems) using the Power SYBR Green PCR Master Mix (Applied Biosystems) and the following specific forward and reverse primers: Agfg1: 5’-CCTGTTGGGAGAGTCTGCAC-3’ and 5’-<colcnt=3> ACCTACAACTGGGGACTGACT-3’; Cd53: 5’-TGCAGATGTTCAGGGTTGCTA and 5’-AAAGGACATTCCCAGCACCT-3’; Cebpb: 5’-CCGGATCAAACGTGGCTGA-3’ and 5’-GATTACTCAGGGCCCGGCTG-3’; Gata3: 5’-TATCCGCTGACGGAAGAGGT-3’ and 5’-CATACCTGGCTCCCGTGG-3’; Gna13 5’-ATCAAAGGTATGAGGGTGCTGG-3’ and 5’-CCACTGTCCTCCCATAAGGC-3’; Hif1a: 5’-ATGGCCCAGTGAGAAAAGGG-3’ and 5’-AGTGAAGCACCTTCCACGTT-3’; Sidt1: 5’-GTCCTCGGAGTGGTGTTTGG-3’ and 5’-ACGGCCCATGTAGTAGATTTGG-3’; Smpdl3a: 5’-AGCTGTGGGGCAGTTTTGG-3’ and 5’- CACACACCTTGGTACGGTCA-3’; Ubap2l: 5’-TTCATTGGGGTTGAGGGGTC-3’ and 5’- TCCATGCACCTGGATGTATCA-3’; Ctla2a: 5’-TCAATTTAGTGACTTGACTCCAGA-3’ and 5’- GGAGCCATTTCTCCTCTATTCAGT-3’; Vim: 5’-AACGAGTACCGGAGACAGGT-3’ and 5’- CAGGGACTCGTTAGTGCCTTT-3’; Gimap4: 5’-CCCAGATTTTCAGGAAGCCGA-3’ and 5’- AAGCTCATGGCTGCTCCTTG-3’; Nfatc1: 5’-GCTGGTCTTCCGAGTTCACA-3’ and 5’- CGCTGGGAACACTCGATAGG-3’; Cxcr3: 5’-GCCATGTACCTTGAGGTTAGTGA-3’ and 5’- ATCGTAGGGAGAGGTGCTGT-3’; Cd200: 5’-TGCCTTACCCTCTATGTACAGC-3’ and 5’-AGTCGCAGAGCAAGTGATGT-3’; Actin b: 5’-CCAACCGTGAAAAGATGACC-3’ and 5’-ACCAGAGGCATACAGGGACA-3’.

For a relative comparison of the gene expression induced by one of the adjuvants in relation to the other, 2^-Δ*CT*^ values were first determined in reference to the *Actb* housekeeping gene of the same sample and then normalized to the average 2^-Δ*CT*^ values obtained for mice immunized with the other adjuvant.

#### Tfh-B cell co-cultures

Groups of 10 C57BL/6 mice were inoculated in the footpads with OVA/IFA or OVA/NP-CpG, as described above. Ten to eleven days later, the popliteal lymph nodes harvested from each group were pooled and the Tfh (CD4^+^CD25^-^CXCR5^+^PD1^+^) and B cells (CD19^+^CD4-) were isolated by FACS. The Tfh and B cells isolated from the same pool were co-cultured in triplicate (30 x 10^3^ and 50 x 10^3^ cells per well, respectively) in round-bottom 96 well plates in complete medium (RPMI 1640 medium (Invitrogen), containing 2 mM L-glutamine, 100 IU/mL penicillin, 100 µg/mL streptomycin, 10% fetal bovine serum, 20 mM HEPES, 50 µM β-mercaptoethanol) and were incubated at 37 °C with 5% CO_2_ without stimulus, with anti-CD3 (clone 145-2C11, eBioscience; 2 μg/mL) + anti-IgM (F(ab’)2-Goat anti-Mouse IgM (Mu chain), eBioscience; 5 μg/mL), or with OVA (EndoFit Ovalbumin, Invivogen; 20 μg/mL). After 5 days, the cells were pelleted by centrifugation and the medium supernatant was collected. The presence of cytokines in the culture medium was tested in multiplex assays (BioLegend’s LEGENDplex bead-based immunoassay, Mix and Match System, according to manufacturers’ instructions; or Eve Technologies service) and the cells were analyzed by flow cytometry for evaluation of isotype-switched B cells (see flow cytometry analysis).

#### RNA-seq processing for bulk sequencing

FACS-sorted cells were collected in DNA LoBind tubes (Eppendorf, #0030108051) containing nuclease-free water (Sigma-Aldrich, #W4502) with 0.2% Triton X-100 (Sigma-Aldrich, #T8787) and RNAse inhibitor at 2 U/mL (RNaseOUT, Invitrogen, #10777019). Cells were frozen in dry ice, kept at −80 °C, and sent to the Genomics Core Facility (GeneCore) at The European Molecular Biology Laboratory (EMBL) for further processing. Briefly, cDNA preparation was done directly from cell lysates according to the protocol described by Picelli et al. for Smart-seq2 (*43*), and sequencing libraries were prepared based on the tagmentation protocol described by Hennig et al. (*44*). All samples were sequenced on NextSeq550 instruments with a high output (**Table S1**).

#### Single-cell library preparation and sequencing

Single-cell libraries of FACS-sorted CXCR5+ Tfh cells from OVA/IFA and OVA/NP-CpG immunizations were generated using the 10x Genomics Chromium Single Cell 5’ V(D)J reagents (10x Genomics; PN- 1000006 and PN-1000020) according to the manufacturer’s protocol. Tfh cells from OVA/NP-CpG and Treg cells from OVA/IFA immunization were loaded together.

#### Bulk RNA-seq Quality Control and differential gene expression analysis

Raw fastq files were aligned against mouse reference genome GRCm38.88 using STAR version 2.5.2a with following parameters: --quantMode TranscriptomeSAM, --seedSearchStartLmax 30 -- outFilterScoreMinOverLread 0 --outFilterMatchNminOverLread 0 --outFilterMatchNmin 30 -- outReadsUnmapped Fastx, for both single end and paired end samples (**Table S1**). The resulting transcriptome aligned bam file was then used as input for quantification using Salmon version 0.8.2 with the following parameters: quant -t -l A. The isoform level counts generated were then used for further analysis.

Tximport version 1.14.2 was used to import and summarize transcript-level estimates for gene-level analysis with countsFromAbundance as the “lengthScaledTPM” option. Transcript to gene file version GRCm38.88 was used, and only protein-coding genes were used for downstream analysis. All genes with Counts Per Million (CPM) of 0.3 or more in at least 27 samples for (Tfh vs. Th) and at least 18 samples (Tfh1 vs. Tfh2) analysis were used. SVAseq from package sva version 3.34.0 was used to estimate surrogate variables, which included batch correction taking into account the strain, response type, and celltype, for all sample analysis and model with strain and response type for Tfh samples only analysis. Limma-Voom package version 3.42.2 was used with “quantile” normalization along with either of the two models (as above) in addition to the respective surrogate variables calculated above to carry out differential expression testing. For Gene Set Enrichment Analysis (GSEA) a reference gmt file of public dataset GSE134153 of only significantly differentially expressed genes with adjust P val < 0.05 was created. A preranked file ordered based on highest to lowest t-statistic with comparison (Tfh vs Th) was created as the input file for the analysis.

#### Logistic Regression analysis to define Tfh1 Tfh2 transcriptome

For this analysis, the Tfh data (27 samples) were used as training datasets, while publicly available datasets (GSE105808, GSE79039, GSE72568) were used as test datasets. Test datasets were first processed, which included alignment and quantification as described above. Genes with CPM of 0.3 or more and expressed in all training and test data were used, resulting in a total of 3571 genes. Svaseq with model “∼ResponseType” was used for batch correction, and batch corrected values were used for downstream analysis. Glmnet version 3.0.1 was used for logistic regression. First, a thousand runs of function cv.glmnet were done for either ElasticNet (alpha=0.5) or Lasso (alpha=1) with “class” as type.measure and “binomial” family on the training dataset. Then the lowest lambda value for option “s” from the runs was chosen for the predictions on the test dataset using function predict.

#### scRNA-seq processing, quality control, and analysis

The scRNA-seq fastq files were processed with CellRanger version 3.0.2 for sequence alignment to GRCm38 (refdata-cellranger-mm10-3.0.0) and per cell quantification of each gene using “chemistry=SC5P-R2”. Seurat version 3.2.3 was used for downstream analysis. Mitochondrial and Ribosomal expression was calculated for each cell using PercentageFeatureSet function. Quality control involved filtering of cells in keeping with the following criteria: cells with a mitochondrial expression of less than 20% and a number of features greater than 200 and less than 4000 were used for downstream analysis. Data were normalized using NormalizeData with default options, and FindVariableFeatures was used with selection.method as vst. Data were then scaled using the ScaleData function regressing out mitochondrial expression, and RunPCA to calculate Principal Components. The top 10 principal components were used to calculate the SNN graph using FindNeighbors function and to calculate UMAP. Data were evaluated for any batch effect. Clustering was done using FindClusters and a resolution of 0.2 to capture a global profile. Based on the expression profile of key follicular (*Cxcr5, Pdcd1, Sh2d1a, Icos*) and regulatory markers (*Il2ra, Foxp3, Ccr7, Klf2*), the clusters were classified as either Tfh or Treg cells. For Tfh2 and Tfh1 analysis, only Tfh cells identified through the above markers were used. These cells were again processed with the initial steps of Normalization and finding Variable Features. CellCycleScoring was done to evaluate the cell cycle state of each cell. Data were scaled, and the percentage of mitochondrial expression was regressed out. The top 20 PCs were used for SNN graph and UMAP calculations using FindNeighbors and RunUMAP, respectively. FindClusters was used for clustering at a resolution of 0.2. A cluster with very low number of genes (200) was excluded. FindAllMarkers was used to find markers for each cluster. Genes with an adjusted p-value of less than 0.05 were considered statistically significant.

#### RNAscope

Popliteal lymph nodes harvested 11 days after immunization of C57BL/c mice were embedded in OCT medium and frozen by immersion in isopentane chilled in liquid nitrogen. 7 µm lymph node sections were stored at −80°C until required. Sections were thawed for 2 min at room temperature, fixed in acetone for 10 min, and air-dried for 2min. They were then rehydrated in PBS for 5 min, incubated in 1.6% PFA for 10 min, washed twice in PBS, and then incubated for 1 h in blocking buffer (0.3% triton, 1% BSA, and 1% mouse serum in PBS). Directly conjugated primary antibodies (**Table S4**) were applied overnight at 4°C, then washed twice with PBS. Sections were then incubated for 30 min in 4% PFA, washed with PBS, and incubated at room temperature for 1 h in ice-cold 70% ethanol. Slides were washed twice with water, then stained for RNAscope probes (Gna13 – Probe Mm-Gna13-O1- C1, Cxcr3 – Probe Mm-Cxcr3-C2, Cebpb – Probe Mm-Cebpb) according to manufacturer instructions. Briefly, sections were washed once in PBS, then incubated for 2 h with the probe of interest in a humidity chamber preheated to 40°C, followed by sequential incubations with four signal amplification ligands at 40°C, each washed twice with 1x wash buffer for 5mins at room temperature. After the final wash, sections were finally mounted in PBS and imaged within 24h.

#### Image analysis

Confocal images of stained sections were acquired using the Zeiss LSM 800 microscope. Following maximum intensity projection and spectral unmixing, images were imported into Imaris for analysis. GL7 staining was used to create a mask for germinal centres, within which CD4^+^ cells were identified as surface objects. The number of RNA particles was counted as spots, then their count per CD4^+^ cell per sample was exported for analysis.

## Acknowledgments

We thank Bruno Silva-Santos, Nuno Morais, Karine Serre, and Jorge Carneiro for helpful discussions. We would also like to thank the Flow Cytometry, Bioimaging, and Animal Facilities of Instituto de Medicina Molecular João Lobo Antunes for their technical support. We thank V. Dangles-Marie, and C. Daviaud, from the mouse facility technicians and C. Guerin, S. Grondin, and A. Viguier flow cytometry core at Institut Curie, as well as the ICGex NGS platform of the Institut Curie (S. Lameiras, S. Baulande, M. Bohec) for technical help with single-cell RNA-seq experiments. This work was supported by ENLIGHT-TEN project, European Union’s Horizon 2020 research and innovation programme under the Marie Sklodowska-Curie grant agreement 675395 to S.K.; UIDB/04046/2020 and UIDP/04046/2020 Centre grants (to BioISI) from the FCT, Portugal to M.G.-C.; Institut National de la Santé et de la Recherche Médicale, Labex DCBIOL (ANR-10-IDEX-0001-02 PSL and ANR-11- LABX0043), SIRIC INCa-DGOS-Inserm_12554 to E.P.; and FAPESP/19906/2014, EXPL/NPN/0082/2019, and LISBOA-01-0145-FEDER-007391, projeto cofinanciado pelo FEDER através POR Lisboa 2020– Programa Operacional Regional de Lisboa, do PORTUGAL 2020, e pela Fundação para a Ciência e a Tecnologia to L.G. A.P.B. was supported by PTDC/CVT-CVT/31840/2017 from the FCT, Portugal.

---

**Supplementary Figure 1:**
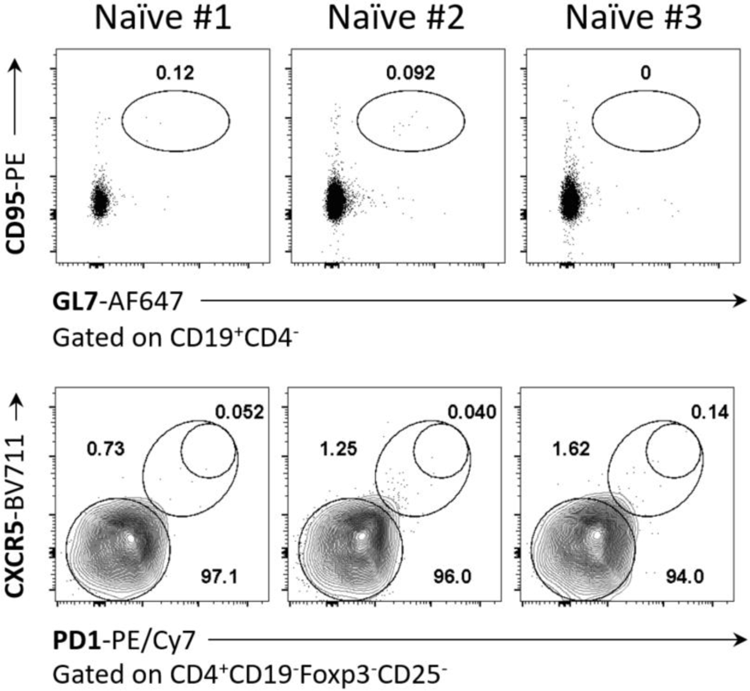
Popliteal lymph nodes from non-immunized mice are devoid of GC and Tfh cells. Representative dotplots of cells from popliteal lymph nodes of three non-immunized C57Bl/6 mice (see Figure 1C as a reference). We could not identify GC B cells (CD95^+^GL7^+^), nor Tfh cells (CXCR5^+^PD-1^+^).

**Supplementary Figure 2:**
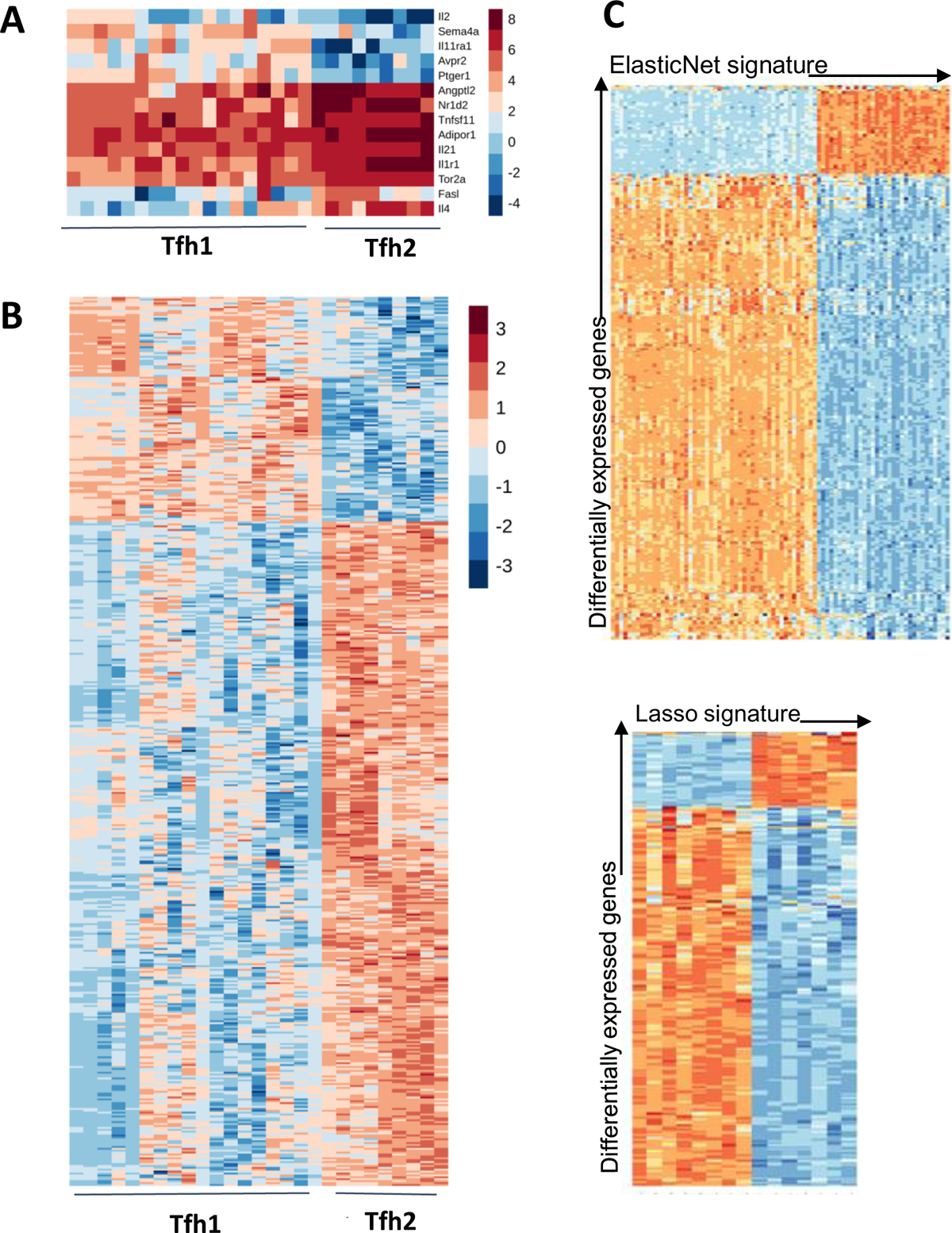
Differential Gene expression of Tfh1 and Tfh2 subsets. (**A)** Unscaled heatmap of genes matched with immune gene list in Tfh2 vs Tfh1 comparison. The plot shows that while *Il21*, *Tnfsf11*, and *Angptl2* are differentially expressed between two subpopulations, these genes are expressed in all Tfh samples consistently. (**B**) Heatmap of significantly differentially expressed genes between Tfh1 and Tfh2 cell subsets from both strains. Genes with adjusted p values < 0.05 were considered significant. Values are scaled for each gene across all samples. The heatmap shows inconsistent patterns of expression from all samples of the same category (Tfh1, Tfh2) (**C**) Correlation of the transcriptome signature from Lasso (top) and ElasticNet (bottom) against the differentially expressed genes.

**Supplementary Figure 3.**
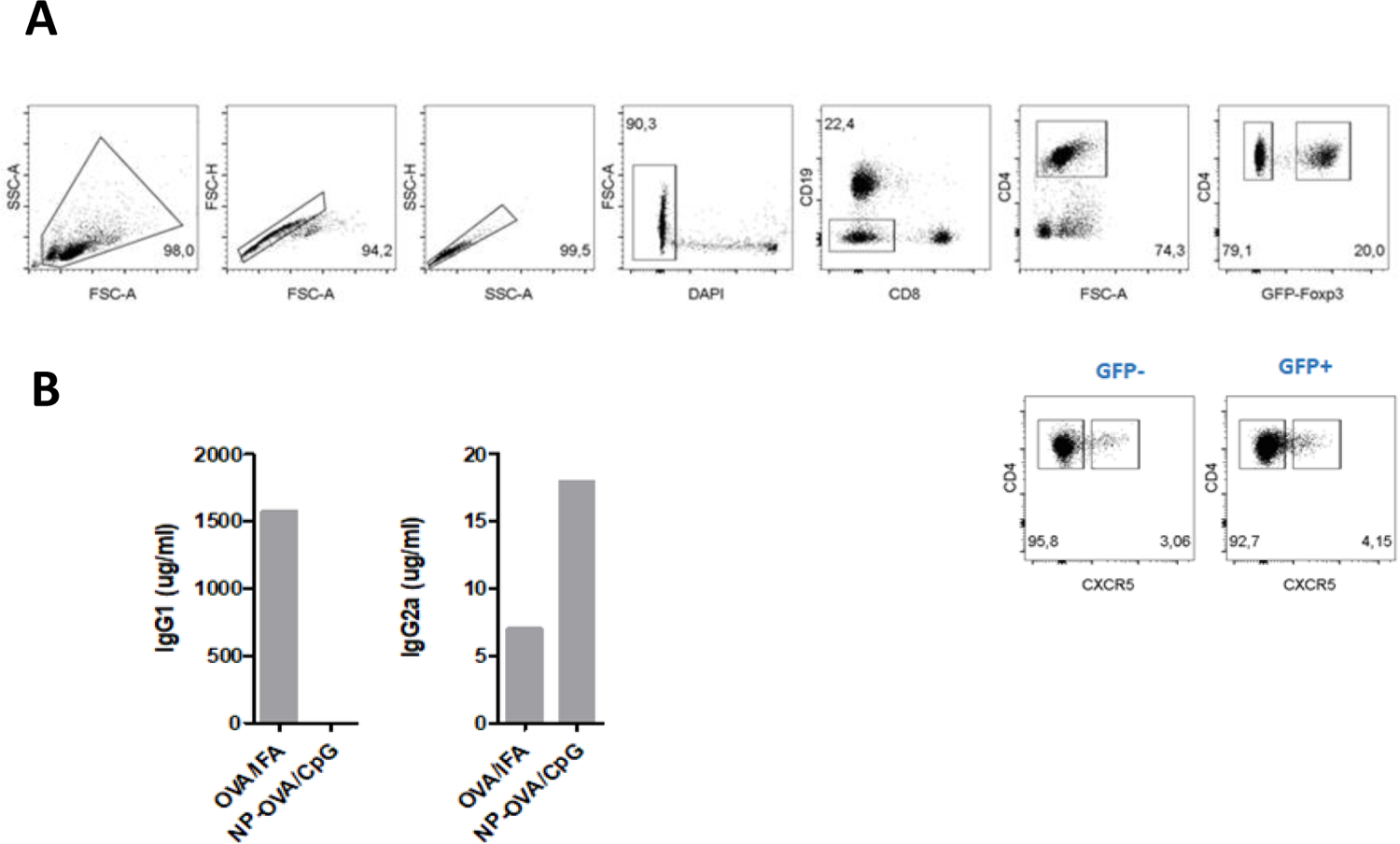
Single cell RNA-seq for Tfh and Treg cells. **(A)** Gating strategy used for cell sorting of Tfh and Treg cells after 11 days of immunization. **(B)** ELISA quantification of OVA IgG1 and IgG2a in the serum of immunized mice.

**Supplementary Figure 4.**
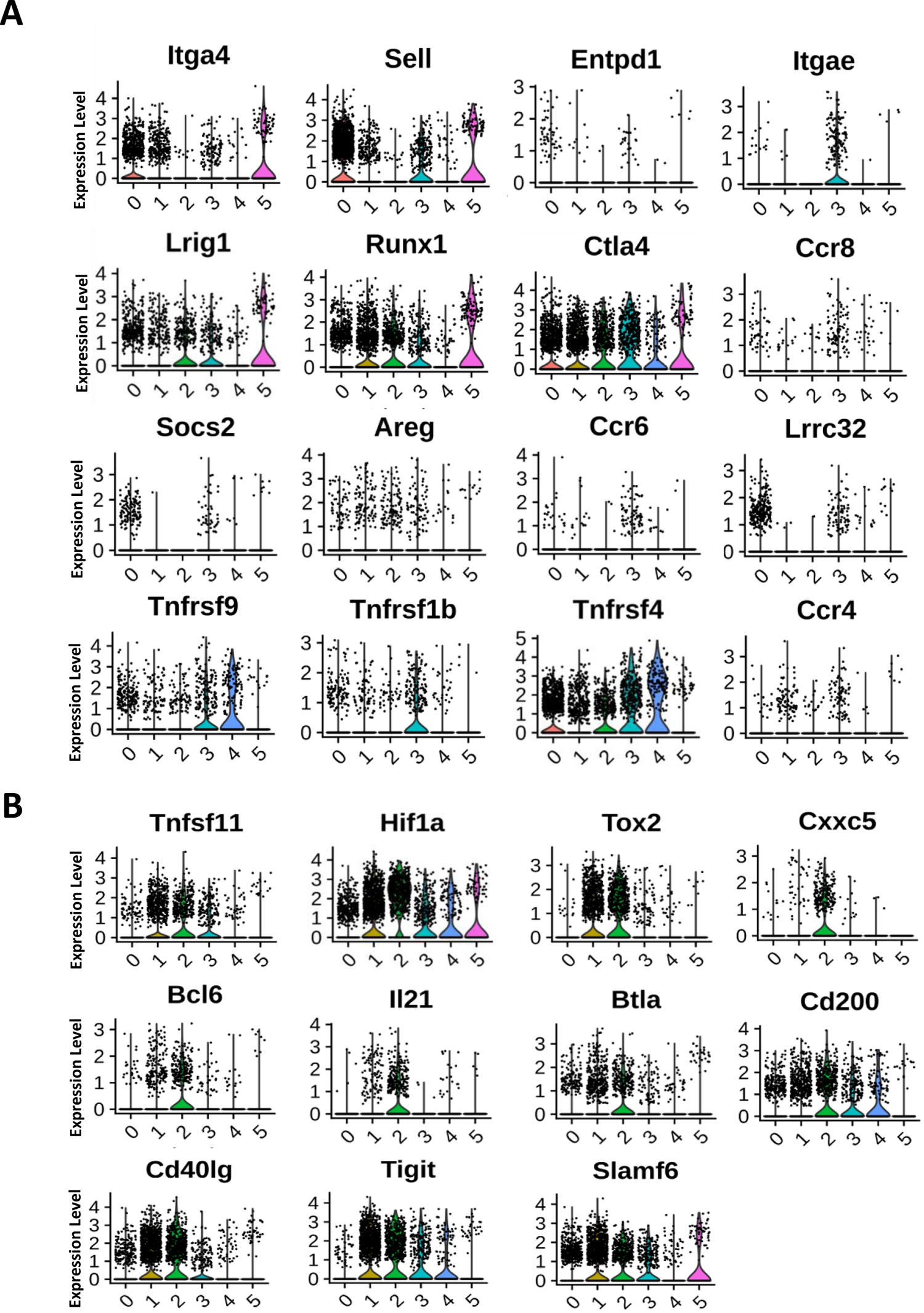
Tfh an Treg markers. Violin plots of a selection of **(A)** Treg and **(B)** Tfh markers describing the heterogeneity of these cells in different clusters.

**Supplementary Figure 5.**
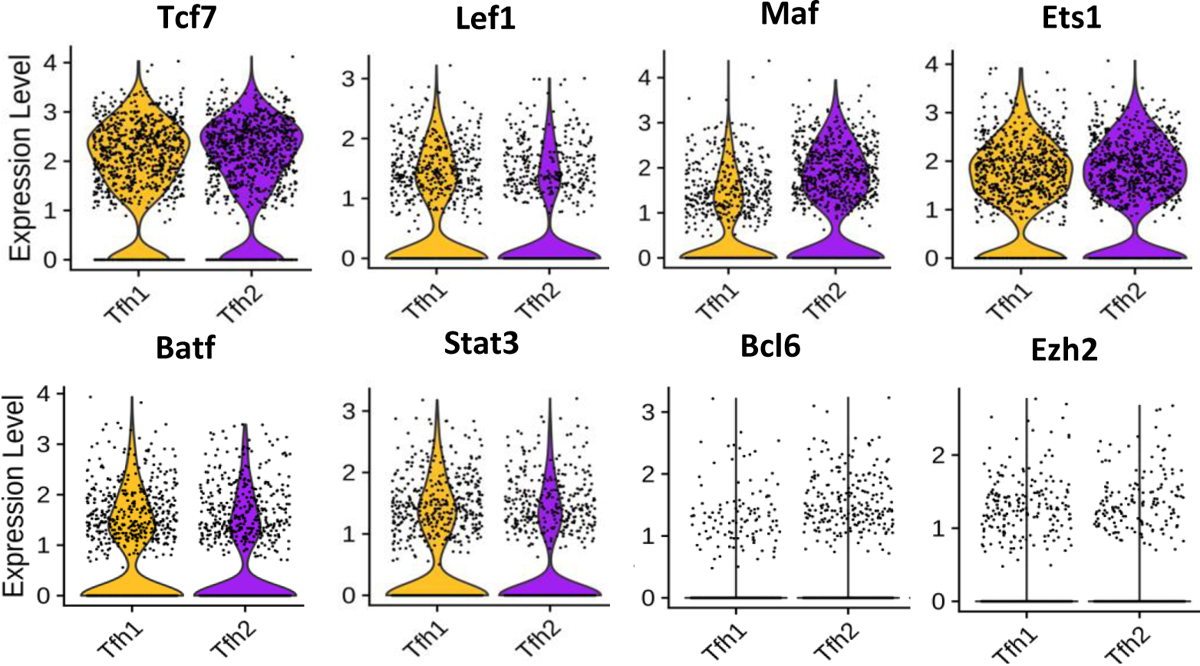
Transcription factors in Tfh1 and Tfh2 cells. Violin plots showing expression of known transcription factors in Tfh phenotype in both Tfh1 and Tfh2 clusters. The plots show comparable expression in both subsets.

**Supplementary Table 1.** Summary of all samples sequenced for each immunization and from the two strains

| Cell Type | Response Type | Mouse Strain | Immunization | Batch | No of Samples | Sequence Type | Read Length | Average no of cells |
| --- | --- | --- | --- | --- | --- | --- | --- | --- |
| Tfh | 1 | C57B6J | CpG | 2 | 5 | Single-end | 75 | 564 |
| Tfh | 1 | C57B6J | NP-CpG | 4 | 4 | Single-end | 85 | 1036 |
| Tfh | 2 | C57B6J | IFA | 2 | 5 | Single-end | 75 | 5000 |
| Th | 1 | C57B6J | CpG | 2 | 5 | Single-end | 75 | 4656 |
| Th | 1 | C57B6J | NP-CpG | 4 | 4 | Single-end | 85 | 1357 |
| Th | 2 | C57B6J | IFA | 2 | 5 | Single-end | 75 | 5000 |
| Tfh | 1 | BALBC | CpG | 1 | 5 | Paired-end | 100 | 70 |
| Tfh | 1 | BALBC | NP-CpG | 3 | 4 | Single-end | 85 | 1011 |
| Tfh | 2 | BALBC | IFA | 1 | 4 | Paired-end | 100 | 350 |
| Th | 1 | BALBC | CpG | 1 | 5 | Paired-end | 100 | 187 |
| Th | 1 | BALBC | NP-CpG | 3 | 4 | Single-end | 85 | 4114 |
| Th | 2 | BALBC | IFA | 1 | 4 | Paired-end | 100 | 275 |

**Supplementary Table 2.**
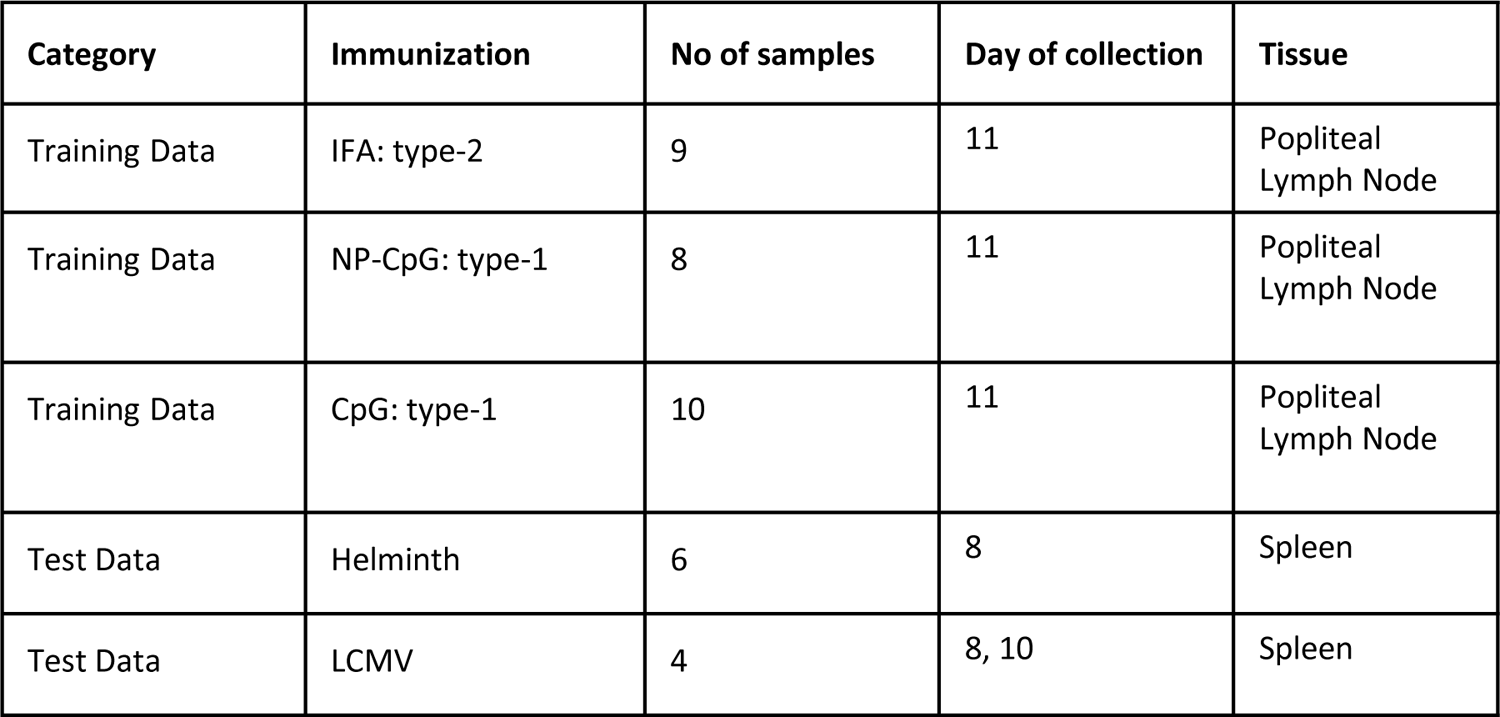
Samples used to generate for transcriptional signature.

**Supplementary Table 3.**
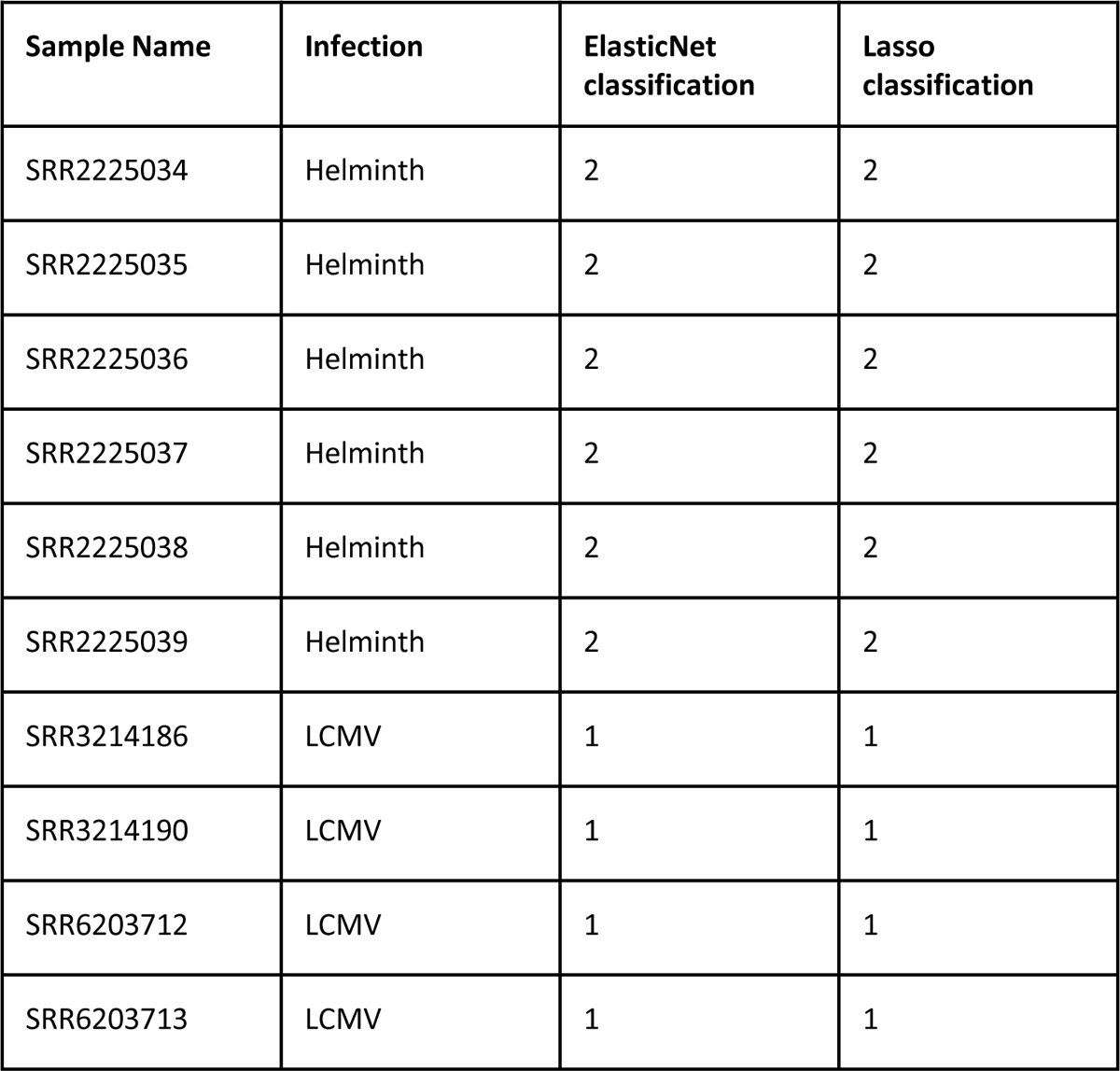
Public datasets used and results of both elasticnet and lasso classifiers in classifying each sample as belonging to either as type 1 or type 2 category.

**Supplementary Table 4.** **Antibodies used for histology**. These antibodies are related to the image in Figure 7C.

| Table S4: Antibody details |  |  |  |  |  |
| --- | --- | --- | --- | --- | --- |
| Target | Clone | Fluorophore | Source | Code | Concentration |
| CD4 | RM4-5 | BV421 | BioLegend | 100543 | 1:100 |
| GL7 | GL7 | AF488 | BioLegend | 144612 | 1:100 |
| IgD | 11-26c.2a | NIR685 | BioLegend | 405749 | 1:200 |

